# The genetic basis for DNA methylation variation across tissues and development

**DOI:** 10.1101/2025.09.15.675351

**Authors:** Jonathan Rosenski, Ofra Sabag, Eitan Marcus, Netanel Loyfer, Yuval Dor, Howard Cedar, Tommy Kaplan

## Abstract

The mechanisms by which genetic variation shapes the epigenome across cell types and developmental stages have remained elusive. Here, we define a unifying developmental framework for DNA methylation programming, grounded in genome-wide methylation and genetic variation data from both mouse and human. In mice, we identify thousands of differentially methylated regions (DMRs) linked to sequence polymorphisms that disrupt transcription factor binding. These DMRs are programmed either during implantation or later in organogenesis, revealing two distinct periods of epigenetic regulation. Extending this logic to humans, we analyze our atlas of over 200 WGBS samples from 39 purified cell types and map 33,574 regions where common SNPs control allele-specific methylation. These include both early-established and cell-type-specific loci, many of which colocalize with eQTLs, enhancers, silencers, and disease-associated variants. Our results uncover a widespread mechanism by which genetic variation influences the regulatory landscape, linking sequence, methylation, and transcription across tissues. This cross-species atlas of sequence-dependent methylation not only clarifies the logic and timing of epigenetic programming, but also provides a foundational resource for deciphering non-coding variants in development, complex disease, and regenerative medicine.

## Main

In mammals, gene expression is governed by both genetic sequences and epigenetic modifications, such as DNA methylation, that instruct the cells of the body when and how to express each individual gene and maintain cell identity through life. DNA methylation patterns are erased in the early embryo and are then re-established during development through highly-programmed interactions between stage-specific and tissue-specific transcription factors that bind local regulatory motifs in the DNA^1^.

There are two critical points when developmental methylation-altering decisions are made. The first is in the embryo at the time of implantation, where almost all genomic CpGs are *de novo* methylated. Nonetheless, some regions, including CpG islands, are actively protected from *de novo* methylation by transcription factor binding to specific motifs, which recruit histone methylases and block DNA methylases within well-defined domains^2^. Once established, these unmethylated regions are then generally preserved in most cell types throughout development. Further points of decision can take place during organogenesis^3^, differentiation^4^ or even postnatally^5,6^, where selected regions undergo tissue-specific demethylation through local transcription factor recognition of specific motifs. While these regions are then maintained unmethylated in the target tissue, they still remain methylated in most other cell types of the body^7^. It is these programmed methylation profiles that actually define stable cell identity^8^.

Analysis of DNA methylation patterns in the mouse has revealed a large degree of variation between different strains used in the laboratory^9^. These differences do not appear to be the result of “spontaneous” methylation changes but rather, are almost always associated with nearby sequence polymorphisms generated during the process of evolution^10^. In this sense, these epigenetic variations are actually due to genetic differences that alter the motif sequences of key transcription factors necessary for setting up local DNA methylation patterns^11^.

In this study, we aim to understand how genetic variation influences DNA methylation across development and in different cell types. For this, we generated a multi-tissue dataset of reduced-representation and whole-genome bisulfite sequencing (RRBS and WGBS) from two mouse strains. By characterizing the molecular basis of genetic drivers that affect the timing and specificity of methylation determination, we deciphered the underlying code of methylation programming during development.

Building on our mouse model findings, we then examined DNA methylation variation in humans, using a comprehensive whole-genome atlas of DNA methylation we recently published, including >200 human samples from 39 primary cell types purified to homogeneity^12^. Taking advantage of the fact that this atlas is derived from non-related outbred individuals, we were able to use the same basic principles observed in the mouse to map the genetic variants and factors that govern the DNA methylation programming network in humans.

Previous studies have already shown significant associations between genetic variation and DNA methylation levels (meQTLs), as well as their relation to complex traits^13–19^. These were mostly limited to blood DNA or used DNA methylation arrays, which only cover ∼1-3% of methylation sites. Here, using deep whole-genome sequencing in purified cells, we were able to capture both genetic and epigenetic information at the single-molecule level, and identify tens of thousands of regions where methylation is determined by genetic variation, which we further analyzed from a developmental perspective - as in the mouse data. To further explore the regulatory role of DNA methylation, we integrated our findings with eQTLs and gene expression data (GTEx), uncovering thousands of genomic regions that function either as enhancers or silencers of gene expression.

This comprehensive catalog of cell-type-specific sequence-dependent methylation elucidates the regulatory mechanisms underlying cellular differentiation, gene expression and phenotype; furthers our understanding of heritable disease and will provide a vast resource for understanding the genetic and epigenetic foundations of gene expression in health and disease.

## Results

During development, there are two major stages where sequence alterations may influence the establishment of DNA methylation patterns. The first decision point is at the time of embryo implantation (E6.5 in mouse), when CpG islands are chosen to be protected from genome-wide *de novo* methylation activity. The second critical event involves the demethylation of selected regulatory regions important for tissue formation and maturation. Motivated by these developmental insights, we investigate the effect of sequence variation on DNA methylation across various tissues in two inbred mouse strains.

We isolated DNA from four distinct tissues (cerebellum, colon, fat, liver) of two mouse strains, C57BL/6 and 129X1/Sv, and performed RRBS to analyze DNA methylation patterns. We identified regions ubiquitously undermethylated in C57BL/6 and methylated in 129X1/Sv across all cell types (Supplementary Fig. 1a), enriched for CpG islands (permutation test p<1e-268). To understand this phenomenon from a developmental perspective, we compared the methylation patterns in the soma to early development (E10.5, C57BL/6). Indeed, these regions are mostly unmethylated at this stage, suggesting the methylation-altering decision was already made early in development, protecting them from *de novo* methylation (Supplementary Fig. 1a).

Similarly, regions ubiquitously unmethylated in 129X1/Sv but methylated in C57BL/6 in all tissue types, were found to be already methylated at the time of implantation in C57BL/6 (Supplementary Fig. 1b). In addition to these ubiquitous DMRs, we also detect genomic regions that show tissue-specific demethylation in only one of the two strains (Supplementary Fig. 1a-b, Supplementary Data 1).

### Whole-genome methylation analysis

The results reported above were obtained by RRBS analysis, which is limited to a small fraction of the genome. To overcome this limitation, we sought an alternate source of data and turned to a recent study mapping genome-wide liver methylation across two pure mouse strains, C57BL/6 and C3H^10^. We analyzed these data using *wgbstools*^20^, and identified 9,095 regions differentially methylated between the two strains in the liver (Supplementary Fig. 2, Supplementary Data 2).

As noted by Grimm and colleagues^10^, a very high percentage of the DMRs were found to be associated with a nearby strain-specific single nucleotide polymorphism (SNP), suggesting that these DMRs are the result of *cis*-acting genetic differences. Presumably, these nucleotide changes alter the methylation programming of associated regulatory regions. As proof of this concept, Grimm et al. carried out crosses between pure C57BL/6 and C3H mice, generating heterozygotes. Liver WGBS data at these 9,095 DMRs from F1 mice, indeed showed an average level of DNA methylation midway between the parents (Supplementary Fig. 2), suggesting sequence-dependent allele-specific methylation (SD-ASM). To contextualize the contribution of genetics - compared to development - on DNA methylation, we conducted a Principal Component Analysis (PCA). This showed that tissue development dominates methylome variation (PC1: 86.2%), while genetics (strain) explains less than 5.6% of the variance (Supplementary Figure 3).

### DNA methylation and development

To analyze the DMRs from a developmental perspective, we examined the methylation pattern of these regions in a variety of adult cell types from pure C57BL/6 mice^21^. Focusing on regions undermethylated in C57BL/6 but highly methylated in C3H liver tissue, two developmental categories emerged (Fig. 1a, Supplementary Data 2), similarly to the RRBS data. The first group consisted of regions methylated in C3H livers, but ubiquitously unmethylated in all adult C57BL/6 tissues, as well as in early embryonic cells (E8.5, Fig. 1a, top). In contrast, regions in the second group were found to be differentially methylated specifically in the liver (Fig. 1a, bottom). Consistent with this, E8.5 embryonic DNA was already methylated at these sites, suggesting they undergo demethylation at a later stage, during liver differentiation in C57BL/6 (but not C3H).

**Fig. 1:**
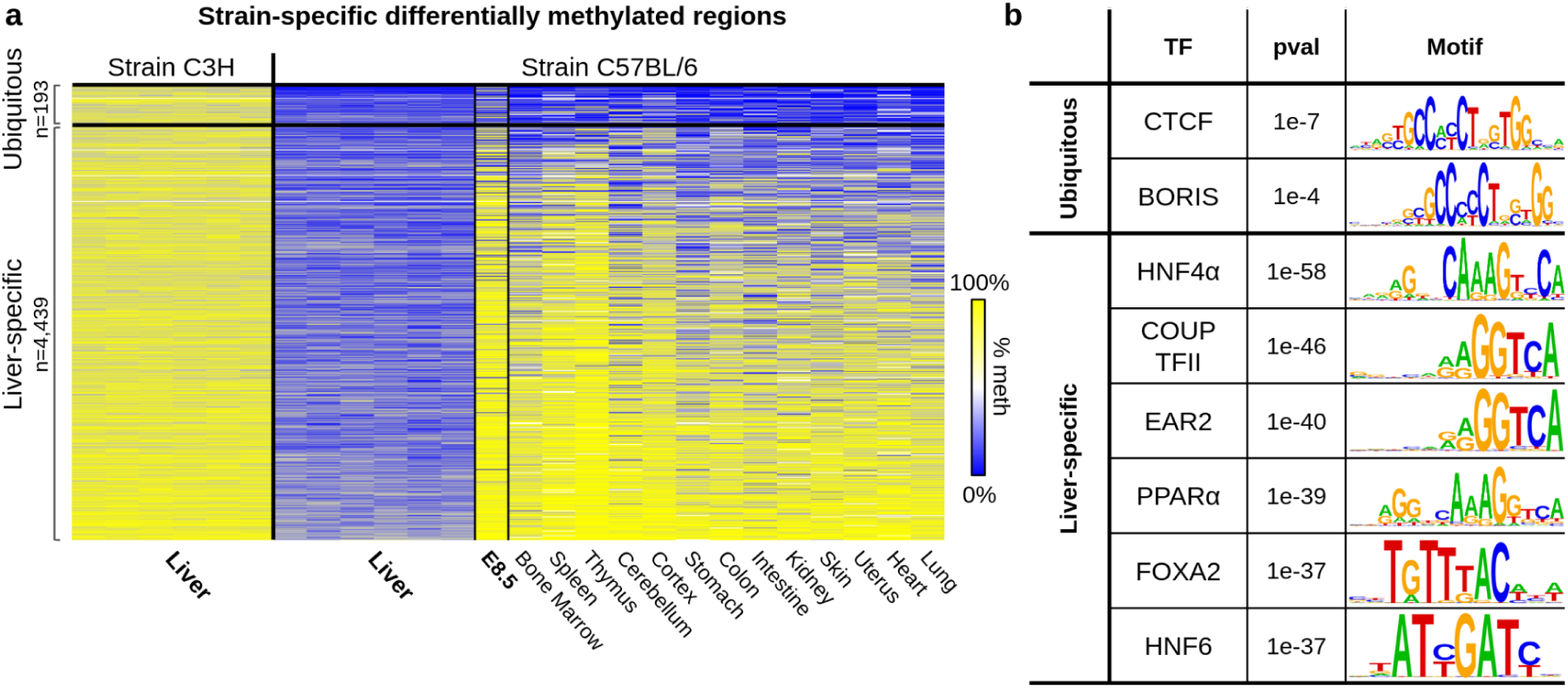
Ubiquitous and tissue-specific DNA methylation differences between two mouse strains. Whole-genome bisulfite sequencing (WGBS) data derived from livers of C3H, and from livers and other tissues of C57BL/6 mice. **a,** Heatmap of statistically significant regions (n=4,632) undermethylated in the livers of C57BL/6 compared to C3H. Examination of these regions in other C57BL/6 tissues reveals two main categories – ubiquitous (top) and liver-specific (bottom) – consistent with early embryonic (E8.5) methylation in C57BL/6. **b,** Sequence analysis of the two groups identified enrichments for global and liver-specific transcription factor (TF) binding motifs.

### Transcription-factor binding at tissue-specific and ubiquitous DMRs

Tissue-specific variations in DNA methylation must be generated from DNA regions programmed to undergo *de novo* methylation at the time of implantation and then retain this pattern in most – but not all – adult cell types. It is the presence or absence of key tissue-specific proteins and their DNA binding sites that direct the DNA demethylation machinery, thereby determining the final methylation state in each tissue/strain.

In keeping with this model, motif analysis of genomic regions specifically unmethylated in C57BL/6 liver revealed highly significant enrichment for key hepatocyte-specific factor families, including HNF, PPAR and FOX (Fig. 1b and Supplementary Table 1). These sites (e.g. FOX) have an intact motif in C57BL/6 DNA, while strain-specific SNPs eliminate TF binding in C3H (Supplementary Fig. 4), thereby preventing demethylation during liver development. Conversely, sequence analysis of the ubiquitous DMRs revealed a different set of enriched motifs, including embryo-specific transcription factors, such as CTCF (Fig. 1b and Supplementary Table 2). A similar number of opposite DMRs, hypermethylated in C57BL/6 but unmethylated in C3H (Supplementary Fig 5a and Supplementary Data 2), was identified, also showing a significant enrichment for liver-specific transcription factors in tissue-specific DMRs (FOX and HNF family motifs), but not among the ubiquitous regions (Supplementary Fig. 5b).

This analysis provides genetic evidence that key tissue-specific “pioneer” factors are involved in mediating the initial demethylation of regulatory regions during organogenesis. Taken together, these descriptive studies of strain-specific DNA methylation differences serve as evolutionary evidence for identifying which regulatory factors play a role in establishing DNA methylation during development.

### Strain-specific DMRs and gene expression

To investigate whether strain-specific differentially methylated regions (DMRs) affect gene expression, we first identified the proximal genes for each DMR. Overall, we found 8.7K genes proximal to C57BL/6-unmethylated DMRs and 7.9K genes proximal to C3H-unmethylated DMRs. Despite the distinct genomic positioning of these DMRs, we observed a significant convergence on target genes, with 5,243 genes (45.6%) common to both strains (nearly twice the amount expected by chance, p<1e−320). Using liver RNA-seq data from C57BL/6 and C3H strains^22^, we found that 1,578 out of 2,190 differentially expressed (DE) genes (72%) are proximal to strain-specific DMRs (p<1e−135). By correlating methylation with expression, we classified these DMRs as either enhancers (negative correlation; ∼4.5K regions) or silencers (positive correlation; ∼3.6K regions, Supplementary Figure 6 and Supplementary Data 3).

### SNP-associated DNA methylation changes in humans

It is well established that much of the variability in human DNA methylation, within one cell type, is associated with genetic differences^15,23–32^. The above analysis using pure inbred mouse strains for studying DNA methylation variation has clearly enabled a better understanding of the molecular and developmental principles governing this process. Indeed, these same insights also provide a simple guideline for deciphering the more complex mechanisms involved in the generation of epigenetic variation in the human outbred population.

To this end, we analyzed our recent genome-wide human DNA methylation atlas, consisting of >200 deeply sequenced samples from 39 healthy primary cell types, purified to homogeneity^12^. These samples were obtained from 137 genetically unrelated individuals, providing a rich source of genetic variation. To systematically identify regions of sequence-dependent allele-specific methylation (SD-ASM)^33,34^, we used a two-step approach. As in Rosenski et al.^35^, we identified bimodal genomic regions which show a mixture of fully-methylated and fully-unmethylated DNA reads (Fig. 2a).

**Fig. 2:**
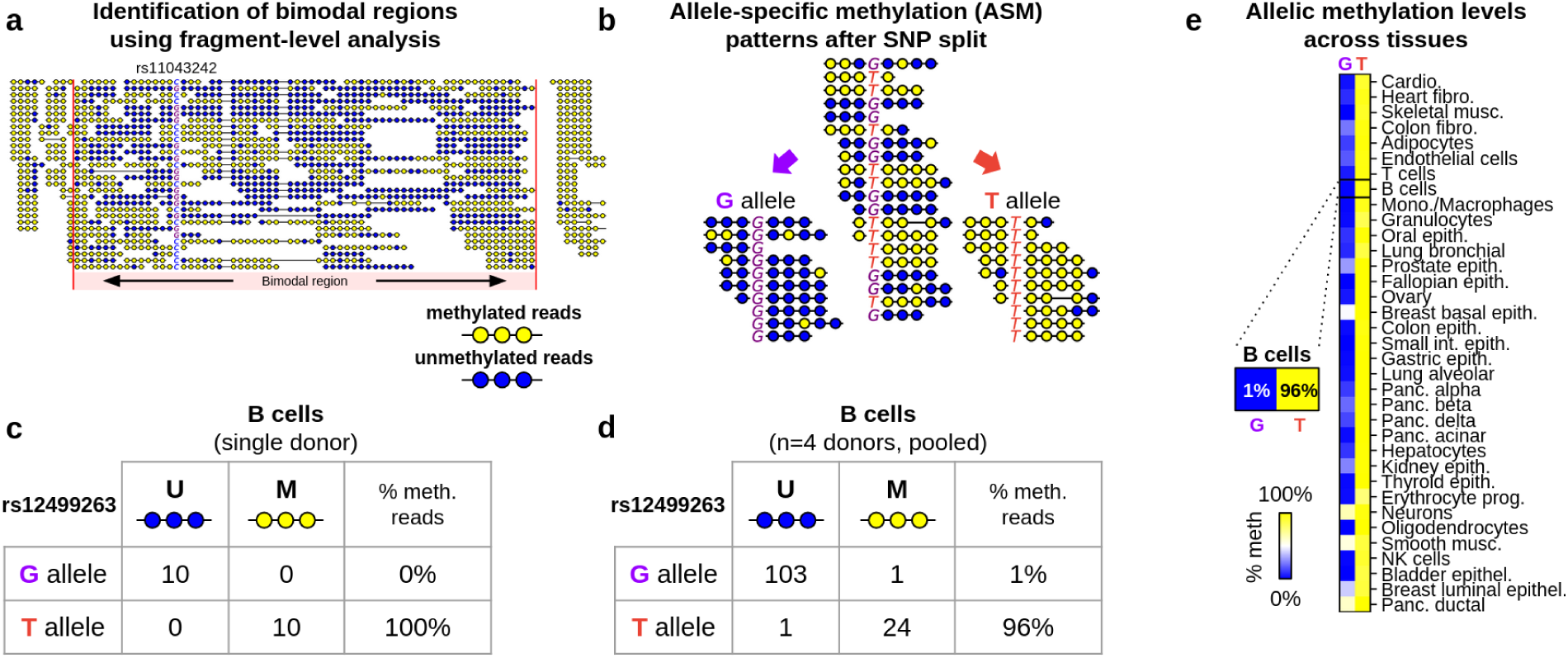
Joint genetic/epigenetic DNA analysis identifies allele-specific methylation regions across cell types and samples. **a,** A computational algorithm identifies bimodal regions^35^, by solely using methylation information of sequenced DNA fragments, of methylomes from 39 purified cell types^12^. Shown, is a bimodal region (chr12:121917805-121919059, hg38, highlighted) where 50% of DNA fragments are methylated (yellow), and 50% are unmethylated (blue), in purified breast basal epithelial cells taken from a single donor, where within this region is a SNP (rs11043242, not used for bimodal region identification) used in subsequent SD-ASM analysis. **b,** Allele-specific DNA methylation shown in fragments from B cells (a single donor) that are segregated by a genetic polymorphism (rs12499263 SNP at chr4:184100474, shown here as reverse complement). Fragments carrying the G allele are mostly unmethylated (blue circles), whereas T allele fragments are mostly methylated (yellow). **c,** Contingency table of alleles by methylation, as shown in b. **d,** Contingency table when n=4 purified B cells samples are pooled across donors. **e,** This allele-specific differential methylation is common across all cell types. Shown is percent of methylated DNA fragments, on the G (left column) and T (right) allele, per cell type.

Bimodally methylated regions represent 5.7% of the human genome and 11% of all CpG sites^35^. Within these regions, we analyzed genetic variation by pooling samples for each cell type, segregating DNA reads by allele at SNP loci, and testing for methylation differences between the reference and alternative alleles (Fig. 2).

Overall, we examined >3 million common SNPs (MAF ≥1%)^36^, and identified 33,574 regions in which methylation is associated with genetic variation in at least one cell type (Supplementary Data 4, 5). We found 166 genomic regions that show statistically significant genetic effects (SD-ASM) in most cell types (termed ubiquitous regions), as well as 19,548 tissue-specific demethylated regions, and 1,173 regions of tissue-specific *de novo* methylation (Supplementary Data 4, 5). The remaining regions did not fall into any single category. In total, SD-ASM regions in humans span >2% of the genome (65 Mb, a total of 1,119,435 CpGs), and are enriched for active regulatory regions including CpG islands (7.5% of regions, n=2,515 out of 33,574, 7.5%, p<1e-260), promoters (7.8%, n=2,609, p<1e-320), and H3K27ac peaks^37^ (53.5%, n=17,974, p<1e-323, Supplementary Figs. 7, 8).

To ensure that SD-ASM is a measure of cis effects rather than population stratification, we determined the genotypes of each individual at 11M SNPs and performed PCA (Methods). Indeed, the top two PCs (ancestry) explained <1% of the SD-ASM regions, suggesting that ancestry is not a major driver of SD-ASM.

The SD-ASM regions tended to group into one of several different categories. In one such regulatory sequence, the reference G allele of rs12499263 (chr4:184100474, hg38) is surrounded by multiple unmethylated neighboring CpGs, whereas the alternative T allele is surrounded by methylated ones. This pattern is ubiquitously found in nearly every sample (Fig. 3a). Furthermore, a fragment-level analysis of individual reads shows all-or-none methylation patterns at the CpGs surrounding this SNP, consistent with the idea of regional epigenetic regulation. This allele-specific differential pattern is observed across all cell types, suggesting that a change in the DNA sequence affected a regulatory element used in the early embryo to protect against local *de novo* methylation, as in the mouse analysis (Fig. 1).

**Fig. 3:**
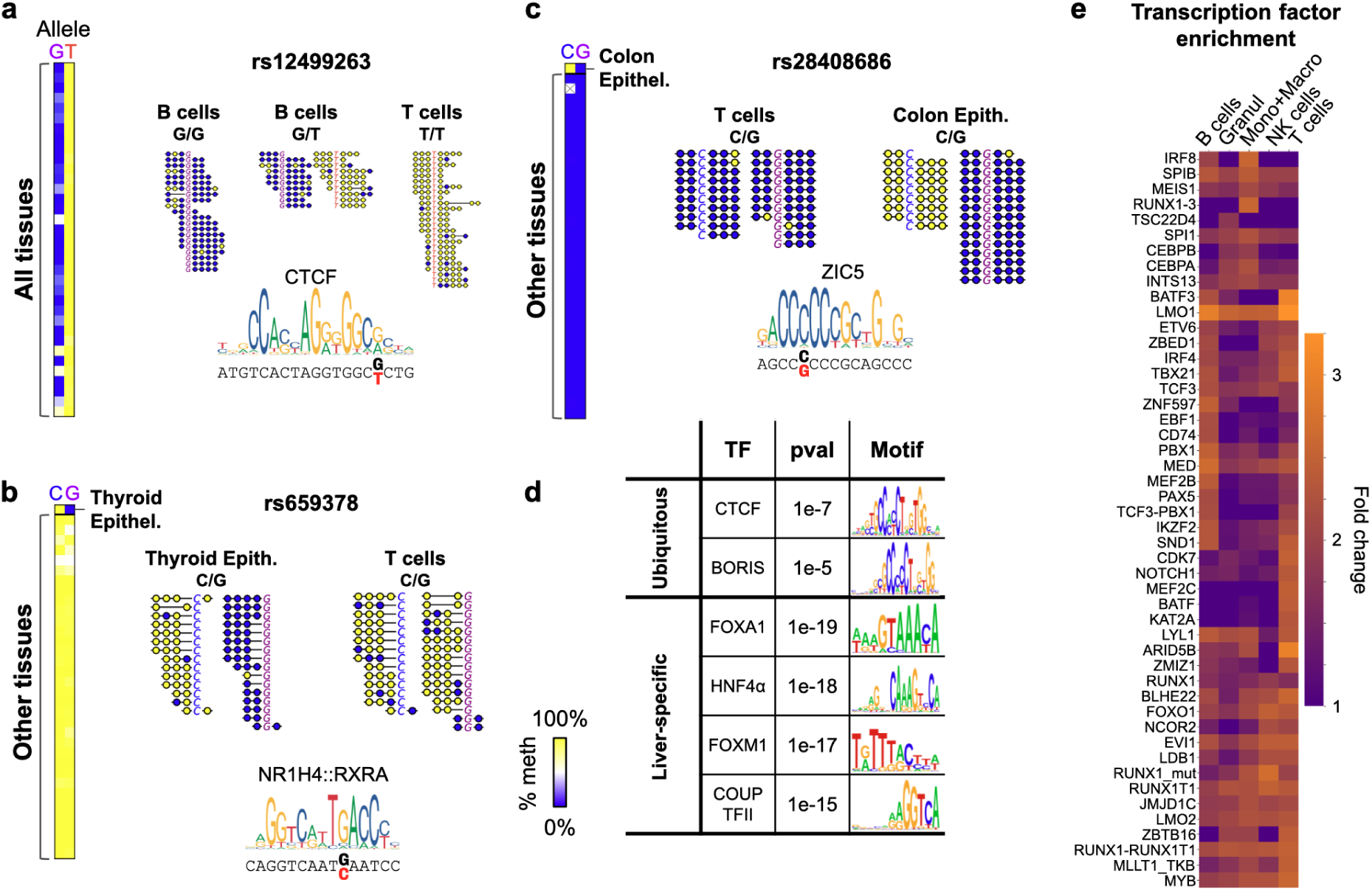
Genetic variation and differential methylation at transcription factor binding sites. **a,** Allelic methylation difference at rs12499263 (chr4:184100474, shown on reverse strand) are ubiquitous across all cell types, where the G allele is undermethylated (blue) and the T allele hypermethylated (yellow). Also shown are sequenced DNA fragments, from three individual donors with G/G, G/T, and T/T genotypes. This SNP overlaps a CTCF binding site, where G-to-T variation hinders binding (e.g. a total of 248 G reads and 6 T reads, in ChIP-seq CTCF data from n=10 donors, ENCODE). **b,** Sequence-dependent allelic differences show G allele-specific hypomethylation in thyroid epithelial cells (top row), but not elsewhere. Also shown are fragments from thyroid and Blood T cells samples, from two heterozygous C/G donors. This SNP overlaps a NR1H4::RXRA motif, where the (unmethylated) G allele is predicted to show higher affinity (rs659378, shown on reverse strand). **c,** Allele-specific *de novo* methylation is shown for the C allele in colon epithelial cells, but not elsewhere. **d,** Motif enrichment analysis for ubiquitous (top), and liver-specific (bottom) allele-specific differentially methylated regions. **e,** Enrichment analysis for *in vivo* transcription factor binding (ReMap)^38^ at tissue-specific SD-ASM regions. Heatmap shows the top 20 factors for each blood cell type, as enrichment fold-change (relative to random), for TFs with FDR<0.05.

Previous work has shown a link between sequence variants and transcription factor occupancy^39^. Indeed, motif analysis showed that the SNP (rs12499263) is located exactly within the binding site of CTCF, an early-embryonic transcription factor known to be required for protection from *de novo* methylation^40^ (Fig. 3a), with predominately higher affinity for unmethylated DNA^11,41,42^. A G-to-T alteration at that position could reduce the binding affinity of CTCF in the early embryo, failing to protect from *de novo* methylation and, therefore, leading to a fully methylated pattern in all tissues of the body (for the T allele). Indeed, analysis of CTCF binding data from 10 donors^43^, revealed a ChIP-seq peak at this specific locus, with a relative 41-fold enrichment of the G allele over T, indicating a preferential binding of CTCF to the G allele, as predicted.

To assess the broader relationship between SD-ASM and transcription factor recruitment, we analyzed all ubiquitous SD-ASM regions intersecting CTCF binding sites^41^. We identified 41 such loci. To validate allelic imbalance at these sites, we used genomic DNA and CTCF ChIP-seq data, and identified 37 unique SNPs that are heterozygous in at least one sample. 30 out of these 37 informative SNPs (81%, including rs12499263) exhibited significant allelic imbalance. This widespread concordance suggests that SD-ASM frequently reflects or facilitates differential chromatin architecture across individuals.

In addition, we identified 25,713 SNPs at 19,548 regions that show cell-type-specific demethylation in one of the two alleles. One such example is a regulatory region on chr18, that undergoes *de novo* methylation at the time of implantation but then is specifically demethylated in the thyroid (Fig. 3b). As in the previous example, a polymorphism (rs659378, chr18:26741849) is located within a transcription factor binding motif, but in this case, a thyroid hormone receptor-related factor is associated with tissue-specific demethylation. Genetic variation at this site, evidently disallows factor binding, thereby preventing the region from undergoing demethylation in the thyroid, but not elsewhere.

In the same manner, but to a lower extent, we identified 1,435 SNPs in 1,173 regions that affect CpG island-like unmethylated regions, which are initially set at the time of implantation, but undergo sequence-dependent *de novo* methylation at specific tissues (Fig. 3c). As in the mouse, ubiquitous SD-ASM in humans is enriched in binding motifs of embryonic factors such as CTCF and BORIS, while cell-type-specific SD-ASM regions are enriched for specialized motifs, e.g. FOX and HNF family factors at hepatocyte-specific SD-ASM regions (Fig. 3d Supplementary Tables 3-4 and Supplementary Data 6).

In addition to cell-type-specific examples, this same process may also take place in multiple cell types, all driven by the same common factor, often because they share a developmental lineage. Supplementary Fig. 9 shows few such examples throughout pancreas development. In blood, Fig. 3e enumerates the most enriched transcription factors, based on ReMap peaks^38^, for regions showing specific sequence-dependent demethylation. While some transcription factors are unique to one cell type (e.g., BATF in T cells), others are enriched across several (or all) blood cell types (e.g., IRF4 in T and B cells, LMO1 in all blood cell types). The statistical significance and degree of enrichment for each transcription factor across all cell types are shown in Supplementary Data 6. By delineating the transcription factors enriched in each specific category (ubiquitous, cell-type-specific demethylation, cell-type-specific *de novo* methylation) we were able to identify putative “pioneer” factors related to that cell type. For example, HNF1a and FOX factors are enriched in hepatocyte-specific demethylated regions, while AATF and NEUROG2 are enriched for neuronal demethylated regions. This multiplex manner of programming generates a complex modular network of hundreds of genomic regions that all undergo coordinated methylation changes (Fig. 4). In this manner, we are able to study which transcription factors affect specific cell types (or groups of cell types). It is worthwhile noting that a number of studies have addressed the connections between SNPs and DNA methylation variation in the human population^14,19,44^. None of these, however, were able to observe the full scope of this system, either because of limited samples and tissues, a lack of cellular purity, or the use of array-based methylation data. Indeed, half of our SD-ASM regions (17,145 out of the 33,574), overlap with previously annotated methylation quantitative trait locus (meQTL) target CpG sites (3.1-fold enrichment, p≤1e-300), whereas the other 16,429 regions were not previously identified^14,16,19^.

**Fig. 4:**
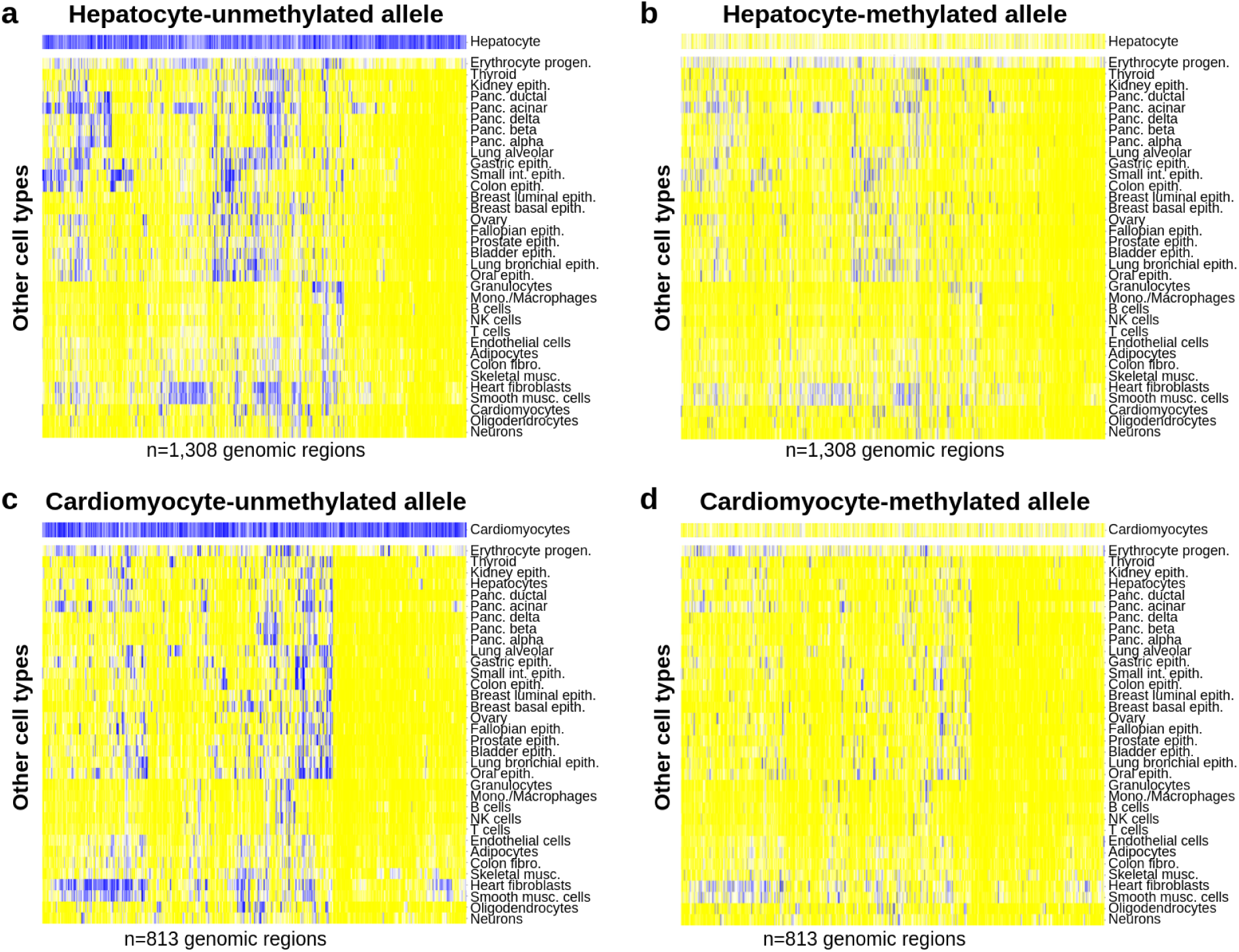
Tissue specificity of allelic demethylation. **a-b,** Hepatocyte-specific sequence-dependent demethylated regions (n=1,308, columns) are shown across all cell types, segregated to the unmethylated allele in hepatocytes (a), and the other allele (b). **c-d,** Same, for n=813 cardiomyocyte-specific allelic loci.

We also analyzed the meQTL effect size of the SNPs that overlap our SD-ASM regions, and report a 2.5- to 3.3-fold increase in beta values, compared to the entire set of previously reported meQTLs^14,19^, presumably because of the increased purity of our samples. Our results thus represent an important advance in understanding the scope and significance of the basic molecular rules that govern DNA methylation programming from the earliest stages of embryogenesis, organogenesis and postnatal development.

### DNA methylation SNPs affect phenotypic variation

With this rich library of regulatory regions, where DNA methylation patterns depend on the genotype of associated SNPs, we then inquired as to whether these epigenetic differences are also associated with the expression of nearby genes, thus contributing to phenotypic variation. To test this possibility, we investigated whether regions of SD-ASM overlap with known expression Quantitative Trait Loci (eQTLs). Using the GTEx dataset^45^, we found 2,618 eQTLs within 500bp of an SD-ASM in the corresponding tissue (1.93-fold enrichment, p<1e-320, Supplementary Data 7), and examined their expression patterns.

For example, the SNP rs12636296 (chr3:125,990,207) clearly shows a widespread effect on the expression of the neighboring gene, *ALG1L* (Fig. 5a). Here, the reference G allele is almost ubiquitously unmethylated and is highly expressed, whereas the alternative T allele is methylated and expressed at significantly lower levels in most tissues, including brain (p≤7e-25), blood cells (p≤3e-22), and heart (p≤3e-19). Another example involves a cell-type-specific SD-ASM region 38Kb upstream of the gene *PCAT18*. Here, both the reference and alternative alleles (rs659378, chr18:26741849) are fully methylated in almost all tissues, but not in thyroid epithelial cells, where only the G allele is unmethylated. Consequently, this allele (G) is significantly upregulated in thyroid cells (p≤8e-180), but not elsewhere (Fig. 5b).

**Fig. 5:**
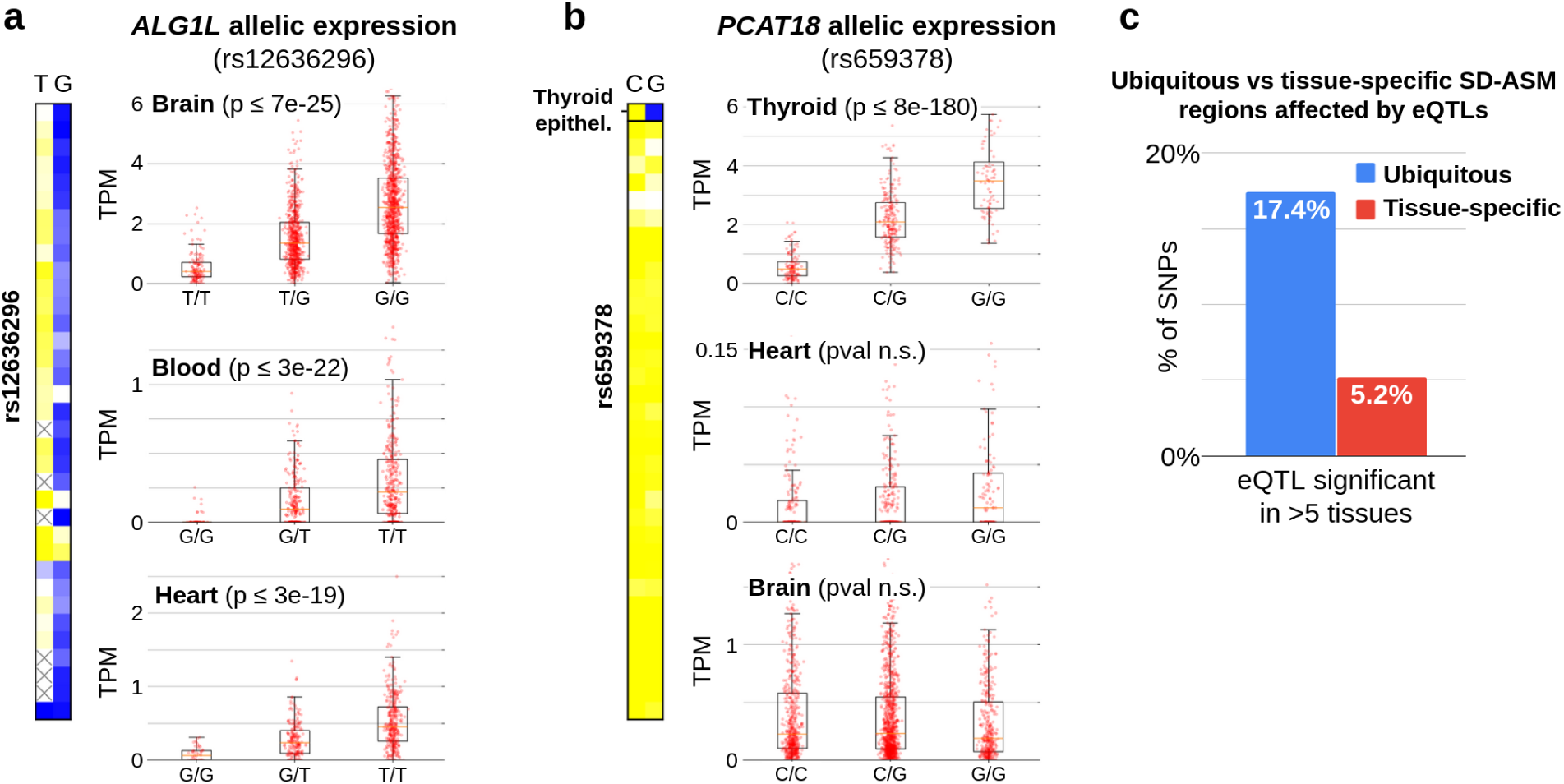
Tissue-specific effects of allelic methylation on gene expression. **a,** Allele-specific methylation at rs12636296 (chr3:125990207, left) shows a ubiquitous pattern, with the T allele methylated and G unmethylated, across all cell types. Accordingly, expression levels of the neighboring gene *ALG1L* (GTEx) differ significantly across donors with different genotypes (at this SNP). Boxplot show the expression (TPM) from brain, blood, and heart samples. **b,** Similar analysis for a cell-type-specific SD-ASM region, which is uniquely unmethylated on the G allele of thyroid epithelial cells (rs659378, chr18:26741849), shows differential expression in the thyroid, but not elsewhere. **c,** Proportion of significant (q<0.05) eQTLs within 100bp of ubiquitous (blue) and tissue-specific (red) SD-ASM loci, showing >5 tissues (p<0.003, Fisher’s exact test).

A global analysis of all SD-ASM regions overlapping known eQTL SNPs revealed significant differences between the ubiquitous and cell-type-specific categories of allele-specific methylation. Genes with cell-type-specific SD-ASM are more likely to show matching expression differences in one tissue, while ubiquitously methylated genes are likely to show an effect in more than 5 tissues (Fig. 5c). In general, it should be noted that in these genetic experiments, all trans-acting factors that control expression are unaffected by the SNP, thereby providing evidence for the role of DNA methylation itself. To formally evaluate whether SD-ASM acts as a regulatory intermediate, rather than a correlated epigenetic phenomenon, we performed two-sample Mendelian Randomization (MR) to test whether SD-ASM acts as a causal intermediate for gene expression^46^. In over 50% of the tested SD-ASM/eQTL overlaps, MR analysis yielded significant results after FDR correction (Supplementary Data 7), supporting the hypothesis that methylation differences are a major driver of differential expression. Additionally, we tested for genetic colocalization of SD-ASM and eQTLs^47^. Among 2,055 significant eQTL/SD-ASM overlaps with sufficient SNP density, 651 demonstrated a high posterior probability of a shared causal variant (posterior probability of colocalization ≥0.8; Supplementary Data 7). We note that our colocalization analysis is limited by read length (assigning p-values to SNPs where reads cover both the SNP and 3 CpGs).

### Disease susceptibility

In the human population there is a great deal of phenotypic variation with regard to disease susceptibility^48–51^. While some of these differences can be explained by variants within the coding regions of genes that take part in trait determination, this mechanism only explains a small percentage of these cases^52,53^ An alternate possibility is that disease susceptibility may result from mutations in regulatory regions that alter their methylation patterns during development and tissue maturation.

Using a computational approach, we analyzed the human SD-ASM library against various datasets and compiled a list of those associated with genes involved in disease. First, we used ClinVar^54^ to associate SD-ASM regions with disease-associated SNPs, and identified 795 SD-ASM SNPs within 500bp of 3,892 pathogenic, or likely pathogenic, hits (p<6e-14, Supplementary Data 8). We then deduced general phenotypes as well as those associated with disease, using the EMBL-EBI GWAS catalog^55^, identifying 1,169 SD-ASM SNPs within 500bp of 889 GWAS hits (p<1e-87, Supplementary Data 9). A colocalization analysis^47^ of SNPs associated with seven hematologic traits vs the SD-ASM signal in six different blood cell types (NK, B, T, Granulocytes, Monocytes, and Macrophages) identified robust colocalized signals in five of these traits; leukocyte quantity, thromboembolism, platelet count, acquired hemolytic anemia, and hematologic disease (Supplementary Data 10).

Finally, to provide a mechanistic understanding of the involvement of DNA methylation variation in human disease, we compared the set of genes whose expression is associated with SD-ASM regions, to the list of known disease-related genes from OMIM^56^, and identified 360 such genes across 16 tissues (Supplementary Data 11). As shown in Fig. 6, rs2980888 is specifically unmethylated in hepatocytes, in the C but not the A, allele. eQTL data from GTEx also link this SNP with *TRIB1AL* expression in the liver. GWAS studies associated this SNP with high-density lipoprotein cholesterol levels (p<6e-170), metabolic syndrome (p<4e-42), cirrhosis (p<7e-9) and various other phenotypes^55^. Thus, while it was previously shown that this SNP predisposes to liver disease via modulation of *TRIB1AL* expression, our findings strongly suggest that the association is mediated via liver-specific effects of the SNP, or an alternative causal SNP, on DNA methylation, which in turn affects liver-specific expression of *TRIB1AL*. Similarly, rs8181996 shows SD-ASM in cardiomyocytes, and is associated with changes in expression levels of *LINC01629* in the heart. This SNP was also found to be associated with atrial fibrillation^55^. Our results thus indicate that the SNP predisposes to heart disease via its effect on DNA methylation, which in turn affects expression of *LINC01629*. We also compared our results to a WGBS-based meQTL dataset^18^, that was found to be enriched in Schizophrenia-associated GWAS variants. Notably, mCpGs that overlap SD-ASM regions showed increased enrichment for Schizophrenia-associated GWAS variants. While general Hippocampus and DLPFC mCpGs regions showed an enrichment (fold change >2), those intersecting with neuronal SD-ASM exhibited much higher enrichment (DLPFC: 12.7-fold, p<1e−45; Hippocampus: 17.5-fold, p<1.7e−43), surpassing the enrichment found in SD-ASM regions alone.

**Fig. 6:**
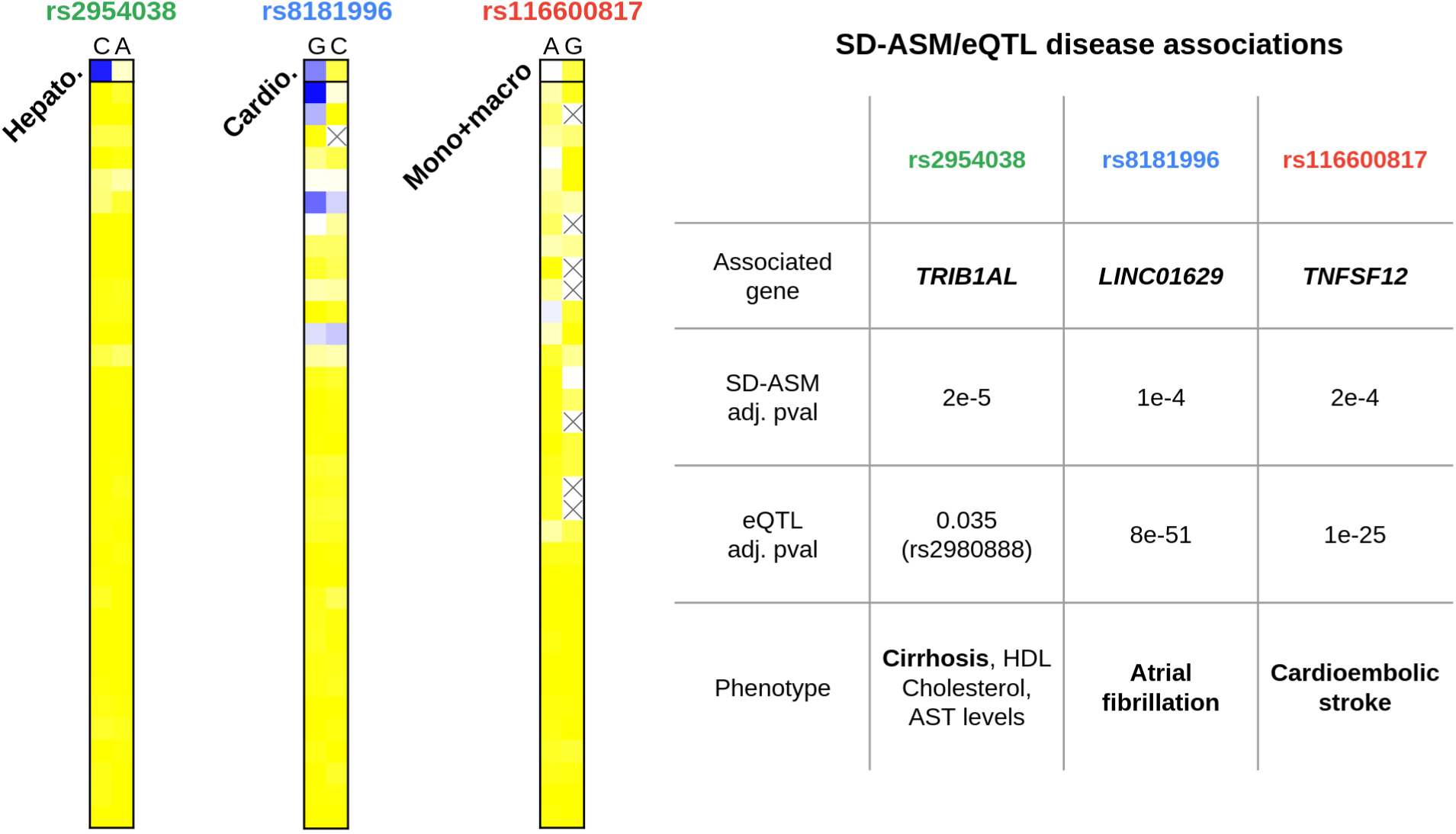
SD-ASM in disease. The SNP rs2954038 is associated with hepatocyte-specific local demethylation of the C allele (chr8:125495066, left), and is associated with increased risk for liver-related phenotypes (NHGRI-EBI GWAS catalog), as well as elevated expression of *TRIB1AL* in the liver (GTEx, via rs2980888). Similarly, rs8181996 (chr14:76961126) is associated with allelic demethylation in cardiomyocytes, as well as differential expression of *LINC01629* in the heart, and increased risk for atrial fibrillation. A third example shows allelic demethylation surrounding rs116600817 (chr17:7555478, A allele) in monocytes, but not in macrophages or other cells. GWAS studies linked this SNP to increased risk for cardioembolic stroke (NHGRI-EBI GWAS catalog). Additional associations are found in Supplementary Data 7-11.

Overall, the significance of these enrichments, as well as the number of associations, suggests that genome-wide programmed regulation of DNA methylation may play a key role in determining inter-individual disease susceptibility and understanding the specific cell types where phenotypes arise.

### Identification of transcriptional silencers and enhancers

Our catalog of SD-ASM, integrated with eQTLs and expression data, provides a unique opportunity to identify how differential methylation is linked with gene expression. Previous examples showed activation properties, where the unmethylated allele is also associated with higher gene expression (Fig. 5). Intriguingly, this is not always the case. Some allele-specific DMRs show the opposite effect, where the unmethylated allele is significantly down-regulated (Fig. 7a). By focusing on SD-ASM/eQTL pairs, per tissue, and examining whether the unmethylated allele is up- or down-regulated, we identified a stringent set of 1,902 statistically significant associations between putative tissue-specific enhancers and their target gene. Similarly, we identified 1,678 associations between putative tissue-specific “silencer” regions and their targets (Fig. 7, Supplementary Data 12-13). Fig. 7a also highlights a few examples, including allele-specific DNA methylation patterns at SD-ASM/eQTL SNPs, and genotype-specific expression data in matched tissues (GTEx). Using the same developmental principles as before, these putative regulatory regions could be further categorized into ubiquitous and cell-type-specific enhancers and silencers (Supplementary Fig. 10).

**Fig. 7:**
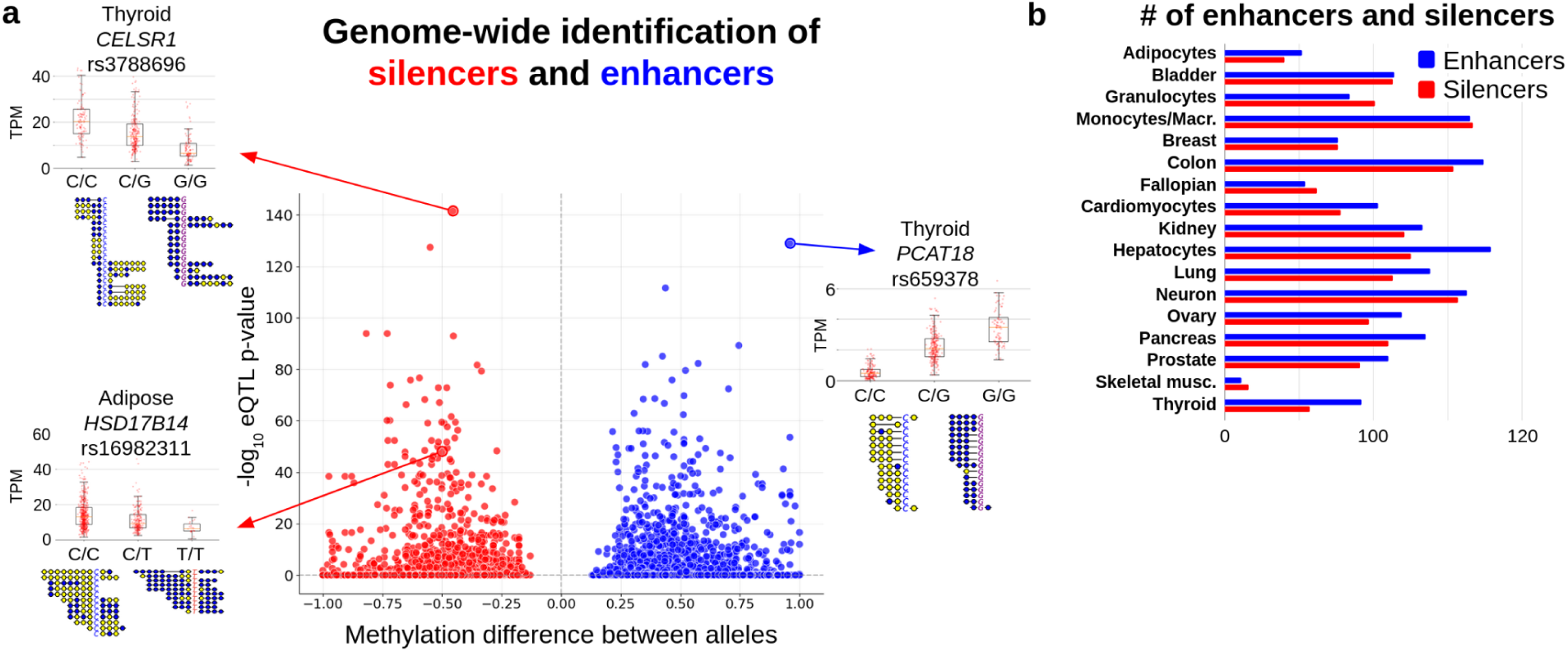
Systematic genome-wide identification of silencers and enhancers. **a,** By comparing the allelic difference in methylation (x-axis) vs. allelic bias in gene expression (GTEx eQTL p-values, y-axis), we identified n=9,189 tissue-specific associations between SD-ASM regions and their target genes. These include 1,902 statistically significant cases of putative “enhancers” (blue), where the unmethylated allele is upregulated in matched tissues, and 1,678 putative “silencers” (red), where it was downregulated. Insets highlight three such cases, including two silencers: *CELSR1* in the thyroid, where gene expression levels (TPM) of individual donors are segregated by genotype (at rs3788696); of *HSD17B14* in Adipose samples (rs16982311) and one enhancer: *PCAT18* in thyroid (rs659378). Below each boxplot. we show a collection of sequenced DNA fragments, split by allele (yellow = methylated CpG, blue = unmethylated CpG). **b,** Bar plots showing the number of cell-type-specific enhancers and silencers, for each cell type.

## Discussion

DNA methylation patterns at genomic regulatory regions are established in a programmed manner as part of embryogenesis. This is accomplished through interactions between local cis-acting DNA motifs and developmental or cell-type-specific transcription factors that can recruit the enzymatic machinery needed to carry out either *de novo* methylation or demethylation. Although this system plays a critical role in setting up stable cellular identity, it has been difficult to pinpoint the factors that mediate these events. In this paper, we have shown that it is possible to use a genetic approach that takes advantage of natural sequence variation between animal strains or within different human individuals, to decipher the basic rules of this epigenetic code.

As a first step, we looked at genome-wide methylation across multiple (bulk) tissues from two pure-bred mouse strains and identified thousands of differentially methylated regions. These are typically associated with polymorphisms that disrupt regulatory motifs, suggesting that specific factors are necessary for setting up proper methylation patterns. From a developmental perspective, these regions fall into two categories. One acts near the time of embryo implantation to locally prevent *de novo* methylation, thereby establishing unmethylated regions that are subsequently maintained throughout life, in all tissues of the body. Genetic variations at those regions could prevent early factors from binding, resulting in ubiquitous hyper-methylation. The second category acts during organogenesis and tissue maturation, causing either demethylation or *de novo* methylation in specific lineages or cell types. In those regions, mutations could presumably prevent binding of tissue-specific factors, thus retaining the embryonic methylation state. Taken together, these observations provide a natural evolutionary tool for deciphering and mapping the factors controlling DNA methylation at regulatory regions^12,57^. While based on whole tissues, the mouse data provides a system where all factors are constant except the genetic make-up of the two inbred strains, with subsequent cross generations providing evidence of real genetically controlled allele-specific methylation. We complement this view by applying the same principles and rationale to human data.

Earlier works studying the genetic variation underlying DNA methylation largely focused on blood and bulk tissues, and often used DNA methylation arrays, restricting their ability to capture the full complexity of epigenetic regulation. Using a deep whole-genome bisulfite sequencing (WGBS) atlas of 39 purified primary human cell types from individual donors, we previously identified hundreds of thousands of genomic regions showing bimodal, allele-specific methylation, with hundreds of them attributed to parental imprinting, often with cell-type-specific differences^35^. Here, we examined the genotype of those bimodal DMRs, across multiple cell types purified from multiple donors, to study their genetic context.

Our results demonstrate that sequence-dependent allele-specific methylation is a widespread mechanism by which genetic variation influences gene regulation. We identified 33,574 such regions, which are highly enriched for CpG islands, promoters, and other regulatory regions. In this study, we leveraged WGBS to identify high-resolution, cell-type-specific associations by analyzing sequencing reads that directly phase genetic variants with methylation patterns. These conclusions are tempered by the modest number of individuals per cell type, limiting the power to analyze associations with ancestry. Nonetheless, it should be noted that a PCA analysis of the SNP genetic profiles suggest that SD-ASM is generally associated with *cis* genetic changes rather than broader ancestral backgrounds. By examining eQTLs and expression data, we identified thousands of SD-ASM/gene pairs, where gene expression significantly varies between the methylated and unmethylated alleles. Importantly, not all unmethylated alleles correlate with up-regulation, as some regions are associated with transcriptional repression, thus highlighting the dual role of DNA methylation in enhancer and silencer function. Strikingly, by linking allele-specific DNA methylation and gene expression, we report a similar number of enhancers and silencers per cell type, suggesting that this unique class of regulation, previously identified using chromatin marks and accessibility changes across different conditions, has been largely overlooked.

These findings underscore the regulatory complexity of SD-ASM and provide a mechanistic link between genetic variation, epigenetic state, and transcriptional output, with implications for understanding both normal development and disease-associated regulatory variation. This approach substantially expands the catalogue of genetically driven methylation changes and enables their classification by developmental timing and cell-type specificity, thereby revealing when and where these changes are established. Notably, half of the SD-ASM regions were not previously detected, and meQTL SNPs that do overlap, show stronger effect size, likely reflecting the increased resolution and purity of our dataset. By integrating genetic, epigenetic, and transcriptomic information within a developmental framework, our study offers a more comprehensive and mechanistic view of how genetic variation shapes DNA methylation across cell types to regulate gene expression and influence phenotypic diversity.

Our findings also have important implications for the epigenetic mechanisms affecting disease susceptibility. While genome-wide association studies (GWAS) have uncovered thousands of disease-linked variants, the functional interpretation of these associations remains limited, particularly for non-coding SNPs. By identifying those SD-ASM regions that overlap pathogenic variants, eQTLs, and known disease genes, we provide evidence that many of these loci act through modulation of DNA methylation at regulatory elements. Importantly, this enables the identification not only of putative causal variants, but also of the relevant cell types in which they act. According to this model, genetic risk, conferred by altered DNA sequence, often acts via its impact on the establishment and maintenance of epigenetic states, offering a mechanistic framework for linking genetic variation to complex disease. As DNA methylation programming is an integral aspect of the body’s long-term response to its environment, our results will also serve as a window for deciphering the hormone-like factors that mediate these effects^57^. Finally, the concepts developed here may provide a new tool for the cell-reprogramming methodologies needed for tissue replacement and regeneration. Using the information obtained here, it may now be possible to identify the key factors required to establish the appropriate cell-type-specific methylation patterns for the successful and stable cell-type conversion and regeneration^8^.

## Methods

### Mice

C57BL/6J and 129X1/SvJ mice were used for all experiments generated in this study. All animal procedures were approved by The Animal Care and Use Committee of The Hebrew University of Jerusalem. Mice were housed and cared for under SPF conditions. Mice were maintained on a 12h light and dark cycle, at 22±2°C, and 55±15% humidity. All experiments used male mice.

### RRBS data

DNA was isolated from fresh mouse tissues (liver and cerebellum) or from isolated cells (white fat cells and colon epithelium cells). All tissues or cells were incubated in lysis buffer (25 mM Tris-HCl (pH 8), 2 mM ethylenediaminetetraacetic acid, 0.2% sodium dodecyl sulfate, 200 mM NaCl) supplemented with 300μg/mL proteinase K (Roche) followed by phenol:chloroform extraction and ethanol precipitation. RRBS libraries were prepared as described^58^ and run on HiSeq 2500 (Illumina) using 100 bp paired-end sequencing.

### Mouse RRBS and WGBS computational analysis

Mouse Reduced RRBS and WGBS data (GSE106379, GSE42836, GSE130735) were aligned to the mouse reference genome (mm10), using biscuit^59^, and methylation analysis was carried out using wgbstools, as described in Loyfer et al.^20^. The ENCODE mm10 blacklist^60^ was used to filter problematic genomic regions in the RRBS data. For WGBS data, CpGs identified as “non-validated” by Grimm et al.^10^ (likely due to mapping errors) were filtered out.

### SD-ASM analysis of inbred mouse strains

To accurately identify strain-specific DMRs and prevent false calls of methylation differences due to CpG-to-non-CpG alterations between strains, we implemented a stringent filtering criterion, including overlap by ≥5 DNA fragments on each strand. Differential methylation calling was run independently on forward and reverse strand reads, retaining only sites showing differential methylation on both.

For RRBS mouse data, wgbstools “find_markers” was used, using a background set of all single CpG, with parameters: min_cov=5, na_rate_tg=0.4, na_rate_bg=0.4, delta_means=0.3, delta_quants=0.3, tg_quant=0.5, bg_quant=0.5, unmeth_quant_thresh=0.5, meth_quant_thresh=0.6, pval=0.01. The “find_markers” command uses methylation blocks consisting of neighboring CpGs. As opposed to smoothing, this segmentation-based method (described in Loyfer et al, Nature, 2023)^12^ groups together consecutive sets of CpGs that tend to change together across various conditions and cell types.

For WGBS mouse data, candidate regions were generated using the wgbstools “segment” command with default parameters, applied to all F0 C3H and C57BL/6 samples. DMRs were subsequently called using “find_markers” with the following parameters: min_cov=10, na_rate_tg=0.2, na_rate_bg=0.2, delta_means=0.3, tg_quant=0.5, bg_quant=0.5, unmeth_quant_thresh=0.5, meth_quant_thresh=0.6, and pval=0.01. In the absence of WGBS data for other cell types or developmental stages in pure C3H mice, we categorized DMRs as ubiquitous or liver-specific based on methylation levels, as in Stadler et al.^61^. Ubiquitous DMRs (Supplementary Fig. 5a-b, top) were defined as those with methylation levels below 10% in the C3H samples. The liver-specific DMRs were defined as those with methylation levels ranging from 20 to 50% in the C3H samples (Supplementary Fig. 5a-b, bottom).

### PCA analysis of mouse WGBS data

Methylation profiles were generated for 5.1M genomic blocks identified using `wgbstools` segment command. Constant blocks (variance < 0.1) were filtered out. PCA was then performed on the remaining (n=12,082) regions to visualize and quantify how genetic strain background and tissue origin contribute to the primary axes of variance across the methylome.

### Mouse RNA-seq analysis

Liver RNA-seq data (TPM) for C57BL/6 and C3H strains (GSE106208) were analyzed to identify strain-specific expression differences. Differential expression (DE) was determined using a two-sided Mann-Whitney U test followed by Benjamini-Hochberg FDR correction (FDR<0.05). Statistically significant genes were then associated with strain-specific DMRs (up to 150 kb). Enrichment of DMRs near DE genes was assessed via Fisher’s exact test against a background of non-DE genes with similar expression levels, selected through a k-nearest-neighbors approach (k=10). DMRs were classified as enhancers or silencers depending on the sign of the correlation between strain-specific methylation and expression (e.g., a silencer would be hypermethylated in the expressed strain).

### Human WGBS analysis

Human WGBS data, sequenced using 150 bp paired-end reads with an average of 984 million pairs per sample, were mapped to the human reference genome (hg38). Data processing and analysis were performed using wgbstools, as detailed in Loyfer et al.^12^. Regions from the ENCODE hg38 blacklist^60^ were excluded from subsequent analyses.

### Identification of bimodal methylation regions

Following Rosenski et al.^35^, bimodal regions were defined as loci showing a mixture of fully-methylated and fully-unmethylated molecules^62–64^, in at least one purified cell type obtained from a single individual, indicative of allele-specific differences at a heterozygous SNP.

Sequenced read pairs were merged and classified based on methylation status. Fragments covering ≥3 CpGs, with an average methylation ≥65% were marked as methylated read pairs (M). Conversely, fragments with ≤35% methylation were marked as unmethylated (U). Fragments with <3 CpGs were excluded, and those with an average methylation between 35% and 65% were classified as mixed (X). Classifying reads as mostly methylated/unmethylated with ≥3 CpGs per read is done to minimize noise. For every genomic CpG position, the proportions of U, X, and M fragments were calculated, and bimodal regions were defined as contiguous regions (≥5 CpGs) with U and M proportions ≥20%, consistent with Rosenski et al.^35^.

A statistical test was devised to differentiate between a null hypothesis (H0) of a single epi-allele (e.g. showing an average methylation of 50%) vs. a mixture model (H1) of two equally probable epi-alleles, as described in Rosenski et al^35^. The identification of bimodal regions depends on the methylation information of sequenced fragments and does not depend on any genetic information. Once bimodal regions were detected in each cell type, a consensus set was determined across all cell types^35^.

### SNPs used for allele-specific methylation analysis

To associate bimodal regions with genotype and sequence-dependent allele-specific methylation (SD-ASM) patterns, we analyzed all 3,046,384 single nucleotide polymorphisms (SNPs) within bimodal regions exhibiting a minor allele frequency (MAF) ≥1% in the GnomAD Ashkenazi Jewish population^36^. To ensure the reliability of our analysis, we applied several stringent filtering criteria. First, we excluded regions from the ENCODE blacklist. To remove variants indicative of poor mapping, we further filtered out any single nucleotide polymorphisms (SNPs) where more than 75% of samples were heterozygous or fewer than 5 samples were homozygous. Samples were classified as heterozygous if each allele had at least five sequencing fragments and comprised between 30% and 70% of the total reads. Finally, we performed an additional validation step by using BLAT with the parameters “-fine -t=dna-minScore=145” to map a 150 bp window around each SNP to the T2T human genome. Any SNP that mapped to two or more genomic loci was removed from the analysis.

To identify sequence-dependent ASM within each cell type containing at least 2 samples, a contingency table was constructed comparing the number of U/M fragments from each individual genotype across all samples of the same cell type. Only SNP-cell type pairs with ≥5 fragments for each allele were retained. Fisher’s exact test was then used to assess the association between allele and methylation patterns, followed by Benjamini-Hochberg False Discovery Rate (FDR) correction^65^.

### Classification of ubiquitous and tissue-specific human SD-ASM regions

For each candidate locus, we classified each cell type individually based on its allelic methylation status at that locus. A given cell type was classified as SD-ASM if it exhibited statistically significant SD-ASM. Alternatively, it was classified as mostly methylated or mostly unmethylated if >70% of its sequencing fragments at that locus were M or U fragments, respectively. Cell types that did not meet these criteria were deemed mixed, and those with insufficient sequencing coverage were designated low coverage. Subsequently, we grouped these loci into three functional categories based on their classification patterns across all cell types. Ubiquitous SD-ASM: Loci classified as SD-ASM in at least 50% of cell types. Tissue-Specific Demethylated: Loci where at least 50% of cell types were classified as “mostly methylated” and fewer than 30% as “mostly unmethylated”. This rule identifies loci with a predominant background of methylation. Tissue-Specific *de novo* methylated: Loci where ≥50% of cell types were classified as “mostly unmethylated” and <30% as “mostly methylated”.

### Enrichment analysis

SD-ASM regions were defined as the bimodal regions containing the SNPs classified as SD-ASM. A bed file was created by adding 100bp to each SNP classified as ASM and subsequently merged, using bedtools merge. These regions were then subjected to enrichment analysis for genomic annotations and motif enrichment using HOMER^66^ with parameters “mm10 homer -bits -size given”. H3K27ac enrichment utilized data from Hou et al.^37^. Data for enrichment analysis presented in Supplementary Fig. 8 were obtained from ChromHMM^67^ predictions on ENCODE data with each annotation merged (bedtools merge) across all samples. For transcription factor binding enrichment analysis of SD-ASM in each cell type (Supplementary Data 6), the Remap dataset^38^ was employed. For all enrichment analyses, statistical significance (p-values) was estimated using a two-tailed permutation test using the enrichment_analysis.py script in Rosenski et al.^35^. Briefly, the number of regions overlapping annotated was compared to the distribution of intersections in 50 randomized selections of the same number of input regions chosen from the background set of all bimodal regions, fitted by a Normal distribution. GWAS data was obtained from the EBI catalog (https://www.ebi.ac.uk/gwas/). The full catalog was used for enrichment analysis with SD-ASM and Supplementary Data 9.

Enrichment of shared genes proximal to mouse and human SD-ASM regions was calculated using a hypergeometric distribution, comparing the observed overlap to the expected intersection of two random gene lists of equivalent size.

### ReMap enrichment analysis

We performed an enrichment analysis to assess the enrichment of cell-type-specific sequence-dependent methylation (SD-ASM) loci with transcription factor (TF) binding sites. For this, we compared the intersection of each set of cell-type-specific methylated and demethylated SD-ASM SNPs with TF peaks annotated in the ReMap dataset (hg38). Specifically, for each list of SD-ASM SNPs, we calculated the overlap of a 200 bp genomic window centered on the SNP with each TF genome annotation from ReMap^38^. This was statistically compared to a background of the same size, which consisted of 200 bp genomic windows centered at SNPs within bimodal regions, which contain similar GC content and are the background from which SD-ASM regions are identified, ensuring strict filtering of false positive enrichment results. The results of this analysis are detailed in Supplementary Data 6.

### CTCF binding

To examine CTCF binding at rs12499263 (chr4:184100174-184100774 hg38; Fig. 4), we used CTCF ChIP-seq data from primary blood cells, across 10 individuals, as listed below^43^. In the main text the reverse strand is used (G/T) while here we refer to the nucleotides on the forward strand (C/A). For 9/10 donors, practically all CTCF ChIP-seq reads contained the C allele, with one donor showing no binding whatsoever, yielding a total of 248 CTCF ChIP-seq reads with the C allele, compared to 6 reads with A, showing a 41-fold enrichment for the C allele. The reported MAF for allele A at rs12499263 varies (q=0.36 in TOPMED, q=0.37 in GnomAD, and q=0.39 in 1,000 Genomes), so a conservative estimate of q=0.3, yields a binomial p-value ≤ 6e-32, computed as 1-binocdf (k=248, n=254, p=0.7). Other (Input) data from the same donors, show A/C heterozygosity and A/A homozygosity at this SNP, further supporting differential binding for the C allele.

| CTCF ChIP-seq<br>ENCODE ID | C allele<br>reads | A allele<br>reads | Other<br>reads | Donor ID | C allele<br>reads<br>(input) | A allele<br>reads<br>(input) | Other<br>reads<br>(input) |
| --- | --- | --- | --- | --- | --- | --- | --- |
| ENCFF165OKY | 0 | 1 | 0 | ENCDO166AGR | 0 | 6 | 0 |
| ENCFF291SPO | 39 | 2 | 0 | ENCDO282XZV | 1 | 1 | 0 |
| ENCFF463NVT | 10 | 1 | 0 | ENCDO383AQS | 3 | 2 | 0 |
| ENCFF467ZUD | 106 | 1 | 0 | ENCDO791UJG | 13 | 5 | 0 |
| ENCFF508JHA | 13 | 0 | 0 | ENCDO781BMZ | 11 | 0 | 0 |
| ENCFF522LXS,<br>ENCFF839WER | 27 | 0 | 0 | ENCDO055UKA | 10 | 0 | 0 |
| ENCFF749JPM | 12 | 1 | 0 | ENCDO195VEA | 1 | 0 | 0 |
| ENCFF749LNR | 24 | 0 | 0 | ENCDO394DKX | 11 | 0 | 0 |
| ENCFF773KHL | 17 | 0 | 0 | ENCDO138EIO | 0 | 0 | 0 |
| ENCFF053RDI | 74 | 0 | 0 | ENCDO439CAZ | 2 | 3 | 0 |

Similarly, to identify allele-biased binding events for CTCF across all ubiquitous regions, we first established genotypes from ENCODE control samples (input DNA). Individual BAM files were processed using bcftools mpileup and bcftools call^68^ to determine biallelic variants at targeted SNP sites. To increase statistical power for identifying heterozygous sites, control BAMs were aggregated using samtools merge before undergoing the same genotyping pipeline.

Allele-biased binding was then evaluated by comparing the allelic read counts in CTCF ChIP-seq data against the baseline allelic ratios observed in the aggregated input controls. Only sites with a minimum coverage of 5 reads in both ChIP and control were tested for allelic imbalance. A two-sided binomial test was performed for each site, with the null hypothesis probability derived from the control allelic ratio using Laplace smoothing (pseudocount=1) to account for low-coverage noise. Enrichment of significant allele-biased events within ubiquitous regions was quantified using Fisher’s exact test.

### Associating SD-ASM with tissue-specific eQTLs

All eQTL data were downloaded from the GTEx Portal (v8). GTEx tissue types were mapped to our purified cell types as follows: Blood (GTEx) to Blood-Mono+Macro (Atlas); Blood Vessel to Vascular endo.; Brain to Neuron; Breast to Breast-Basal-Ep; Colon to Colon-Ep; Heart to Heart-Cardio; Kidney to Kidney-Ep; Liver to Liver-Hep; Lung to Lung-Ep-Alveo; Pancreas to Pancreas-Acinar; Prostate to Prostate-Ep; Small Intestine to Small-Int-Ep; Stomach to Gastric-Ep. GTEx participant genotype information was accessed via dbGaP as VCF files. Statistical significance was estimated using a shuffling test, as described above.

### Comparison to previous meQTL studies

We compare our identified SD-ASM regions to CpG targets of the meQTL studies of Min et al., Hawe et al., and Oliva et al.^14,16,19^. To compare effect size we use the absolute values of the reported beta-values of the meQTLs which intersect our SD-ASM regions compared to the average beta-values of all meQTL per study. Full summary statistics of meQTL/mCpG analysis of Mandell et al.^18^ were obtained with Schizophrenia GWAS summary statistics obtained from Pardiñas et al.^69^. The mCpGs were filtered to have a maximum distance of 300 base pairs from their meQTLs so as to be comparable with the SD-ASM methodology used here.

### Identification of transcriptional silencers/enhancers

To classify SD-ASM/eQTLs as enhancers or silencers, SNPs identified as SD-ASM were initially intersected with all GTEx eQTLs. For each tissue, SD-ASM effects were considered present if the corrected p-value was < 0.05. If, in the same tissue (using the methylation-GTEx mapping defined above), the median expression of the unmethylated allele was higher than that of the methylated allele, the region was classified as an enhancer. Conversely, if the median expression of the methylated allele was higher, it was classified as a silencer.

### Silencers and enhancers found due to linkage disequilibrium

To identify eQTLs in linkage disequilibrium with SD-ASM SNPs, we considered eQTLs of corresponding cell type (using the GTEx to methylation tissue mapping) within a 5000 bp window of the SD-ASM SNP, using bedtools intersect^70^. Each eQTL-SD-ASM pair was then tested for linkage disequilibrium using the LDlink API^71^. We considered only pairs with a significant association (p-value < 0.001) and were classified as being in linkage disequilibrium. Finally, to determine which specific alleles were linked, we analyzed contingency tables of their joint allele frequencies. We applied a stringent filter, requiring that the frequency of an eQTL allele be enriched by at least three-fold on haplotypes carrying one specific SD-ASM allele compared to the other. Only pairs meeting this criterion were classified as in linkage.

### Mendelian Randomization and colocalization

SD-ASM effect sizes were calculated as the log-transformed odds-ratio of methylated (M) to unmethylated (U) fragments across alleles, with a pseudocount of 0.5 to adjust for zero values. Specifically, for allele 1 let 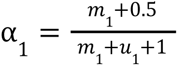 be the proportion of methylated fragments on allele 1 and α_2_ be the proportion of methylated fragments on allele 2. Using a variation of the M-value definition^72^, we define *M*_1_ as 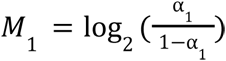, and measure the effect size as β = *M*_1_ − *M*_2_. The variance, σ^2^, of the effect sizes is estimated using Woolf’s formula^73^, and two-sided p-values were derived via a Wald test 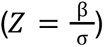.

We then performed Mendelian Randomization (MR) analysis using the TwoSampleMR package^46^ to evaluate the causal relationship between DNA methylation (exposure) and gene expression (outcome). Instrument variables were selected from common SNPs exhibiting significant methylation associations (p<0.05). Standardized effect sizes and standard errors for SD-ASM were calculated and compared to GTEx effect sizes, which were derived directly from published summary statistics.

Statistical colocalization was performed using the coloc package^74^ to determine if SD-ASM/gene expression, and SD-ASM/GWAS signals share a common genetic basis. For each region-gene pair, we examined SNPs for which the effect size was calculated both for methylation and for GWAS/GTEx summary statistics. Variants with high posterior probability (PP.H4≥0.8) were identified as high-confidence colocalized signals.

### Effect of local alleles vs broad genetic profile (ancestry)

The broad genetic profile (ancestry) for each individual was estimated through PCA (PLINK2, MAF ≥ 5% with LD pruning) of genome-wide SNP profiles which were calculated using wgbstools^20^. To differentiate the influence of local genotype from population structure, we employed an allele-level linear regression model where the methylation ratio for reference and alternative alleles served as the dependent variable. The local allelic status and the first two ancestry principal components were standardized and included as independent variables in an Ordinary Least Squares (OLS) model. Standardized slopes and p-values were then calculated to determine the relative contribution of sequence-dependent effects versus global ancestry to the observed methylation variance at each significant site.

## Data Availability

Mouse WGBS data is available from Grimm et al^10^. Mouse RRBS data have been deposited in the Gene Expression Omnibus (GEO) and will be available upon publication. Human WGBS data, raw and processed, are available via EGA (EGAS00001006791) and GEO (accession no. GSE186458). Mouse RNA-seq data was downloaded from GEO (accession no. GSE106208). Any additional information required to reanalyze the data reported in this paper is available from the lead contact upon reasonable request. For GWAS colocalization analysis, we used data downloaded from EBI catalog: hematologic disease: GCST90475821, thromboembolism: GCST90476007, polycythemia vera: GCST90479825, myeloid leukemia: GCST90479832, acquired hemolytic anemias: GCST90479967, platelet count: GCST90480650, white blood cell count: GCST90480724.

## Code Availability

The code will be made available by the date of publication. Additional details and implementation of the bimodal methylation analysis are available at https://github.com/yonniejon/imprint_atlas.

## Acknowledgements

We thank Ben Glaser, Nir Friedman, Yotam Drier, Shai Carmi and members of the Kaplan, Dor, and Cedar labs for insightful discussions, and Ms. Tzippi Jakubowicz for organizing and editing the manuscript for publication. This research was supported by research grants from the Center for Interdisciplinary Data Science Research (T.K., Y.D.), the Howard Jonas Foundation (H.C.), the Israel Science Foundation (grants 1250/18 and 259/23 to T.K.), and the Ministry of Innovation, Science & Technology, Israel (grant 0005421).

## Author Contributions

J.R. and O.S. contributed equally to this work. J.R., O.S., Y.D., H.C. and T.K. conceived the study. J.R., O.S., E.M., N.L. and T.K. performing experiments and analysis. J.R., O.S., Y.D., H.C. and T.K. wrote the manuscript. H.C. and T.K. funded and supervised this study.

