## Supplemental Information (Figures and Tables) for "The genetic basis for DNA methylation variation across tissues and development"

Supplementary Information
Supplementary Figure 1-10
Supplementary Table 1-4

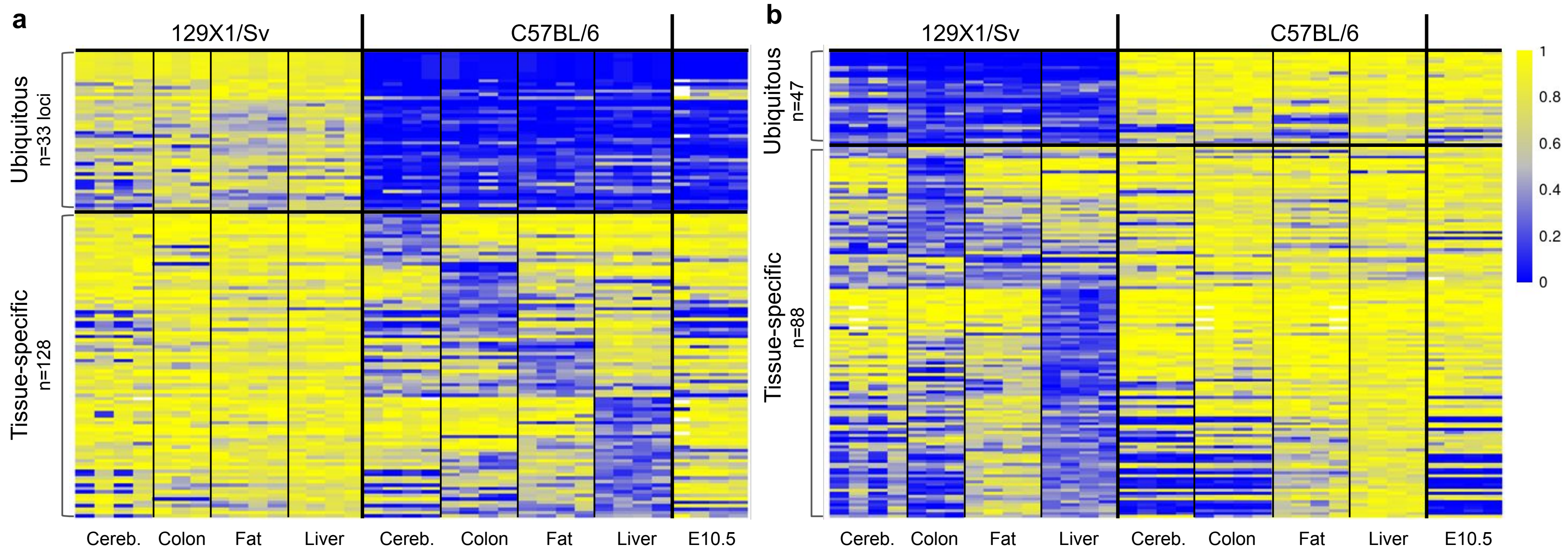


**Supplementary Figure 1. DNA methylation differences observed between two mouse strains.** RRBS data from 4 different isolated tissues (Liver, cerebellum, fat, colon, n=3-4) from inbred C57BL/6 and 129X1/Sv mice. Heatmap of statistically-significant **a,** C57BL/6-specific unmethylated regions and b, 129X1/Sv-specific unmethylated regions. Two main categories (ubiquitous and tissue-specific) could be delineated by comparing these regions across cell types. In addition, methylation levels are shown for the implantation stage in C57BL/6 embryos (E10.5).

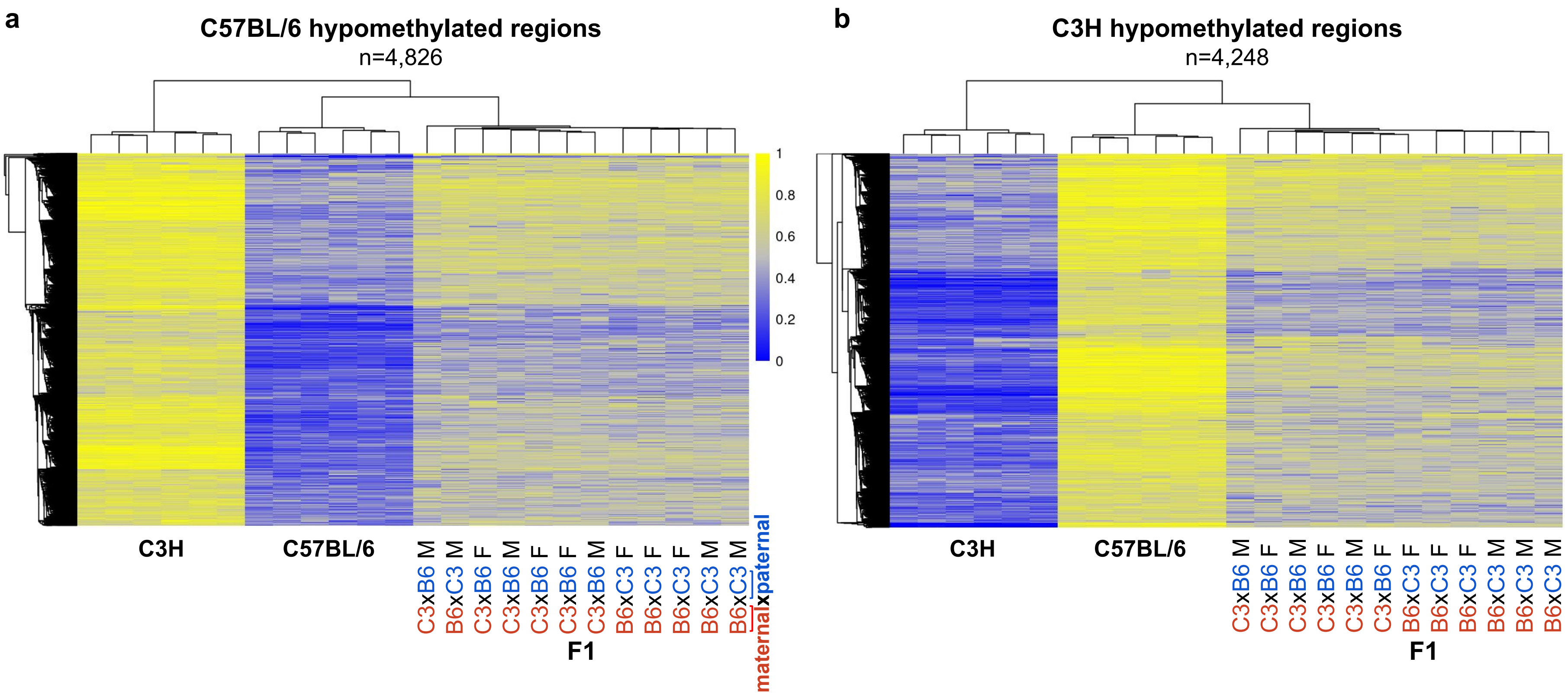


**Supplementary Figure 2. Strain-specific DNA methylation differences are the result of genetic variations.** Heatmap of statistically significant DMRs (n=4,826) unmethylated in C57BL/6 as compared to C3H Liver **a,** or DMRs (n=4,248) unmethylated in C3H as compared to C57BL/6 Liver **b,** These data were also compared to methylation levels in F1 (C3H X C57BL/6) mice.

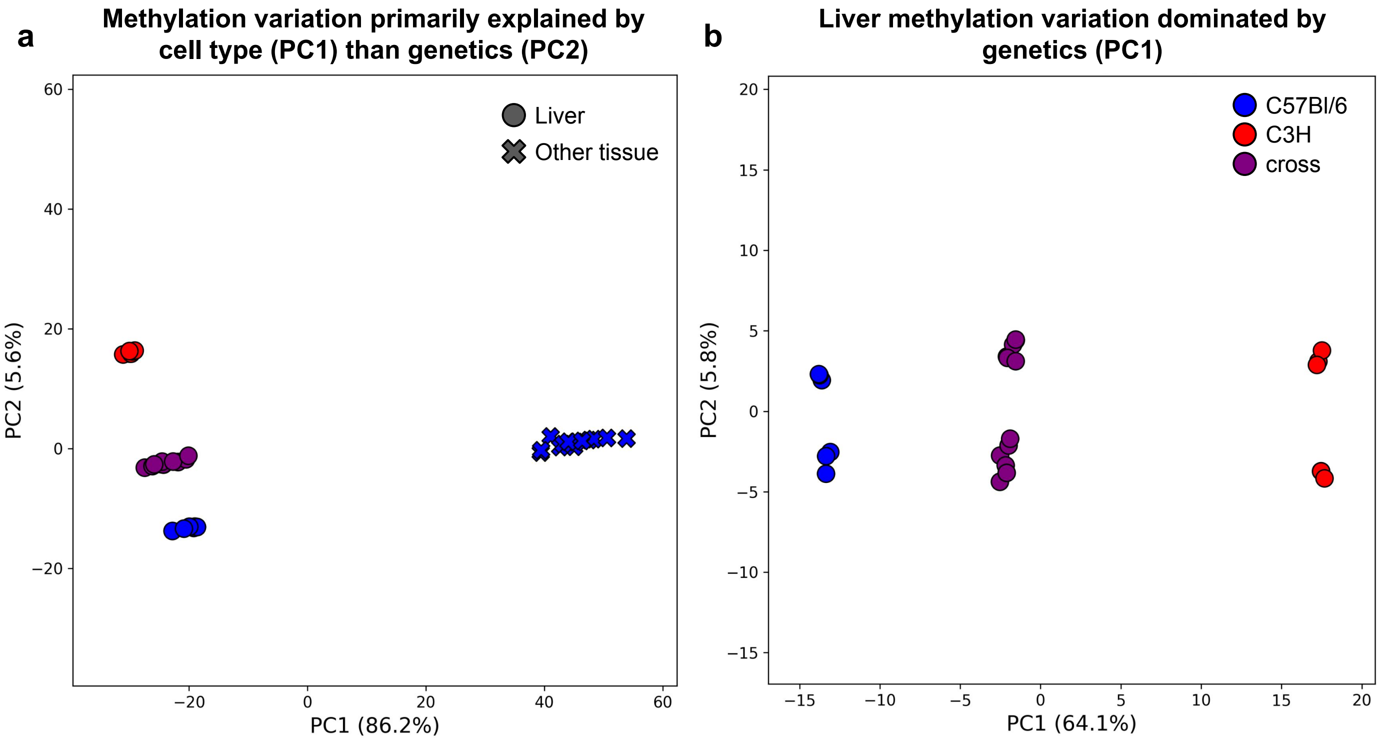


**Supplementary Figure 3. PCA using all liver samples compared to all samples. a,** When only liver samples are used, strain emerges as a top principal component where PC1 clearly distinguishes between the strains and explains >64% of the variance across the methylomes. **b,** When using all samples (not just from the liver) the top principal component clearly separates tissue, explaining > 86% of the variance.

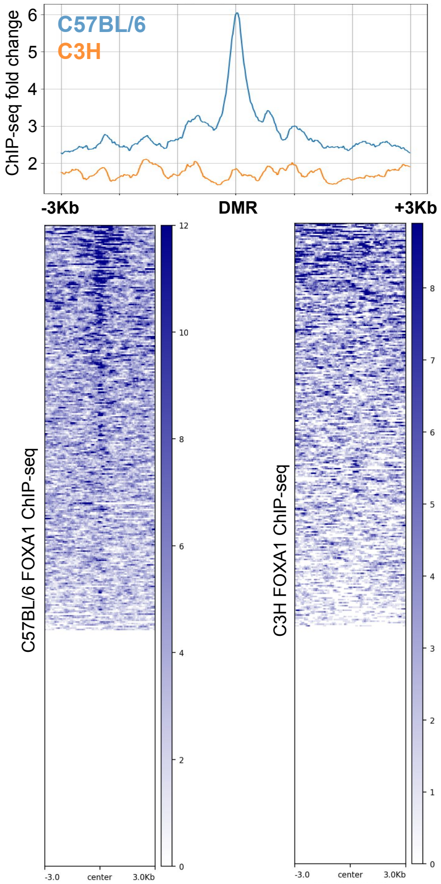


**Supplementary Figure 4. SNPs affect DNA binding activity of FOXA1** Heatmaps of FOXA1 Chip-Seq peaks in C57BL/6 and C3H Liver at hypomethylated regions in C57BL/6 but not in C3 that contain the FOX motif.

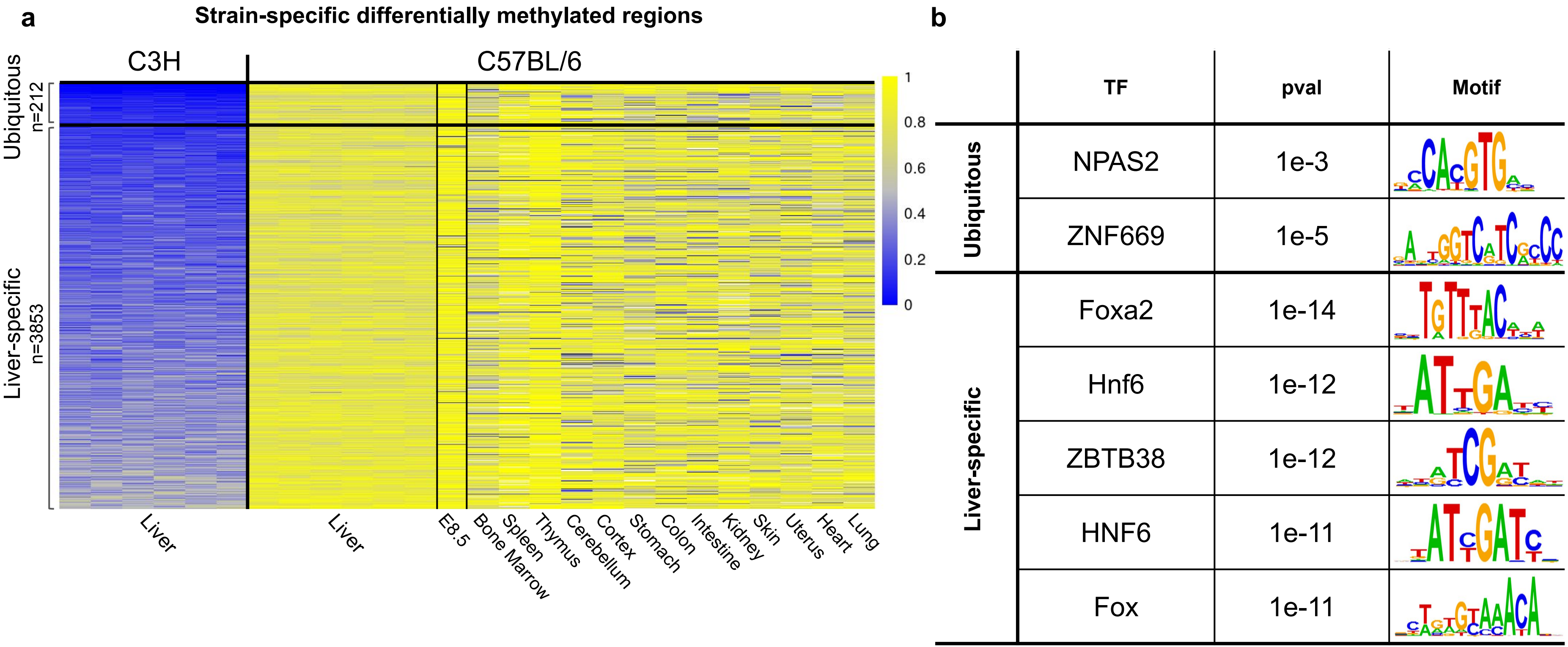


**Supplementary Figure 5. Two main categories of DNA methylation differences observed in mice strains.** Whole-genome bisulfite sequencing (WGBS) data derived from Livers of C3H and C57BL/6 mice. **a**, Heatmap of statistically-significant methylated segments (n=4,065) measured in C3H as compared to C57BL/6 liver. Two main categories (Ubiquitous and Liver specific) could be delineated by dividing the segments into unmethylated (<10%) and low methylated (20-50%) sequences. **b**, Transcription factor (TF) motif analysis (Homer) of these same regions was tested in both the ubiquitous and liver-specific sites.

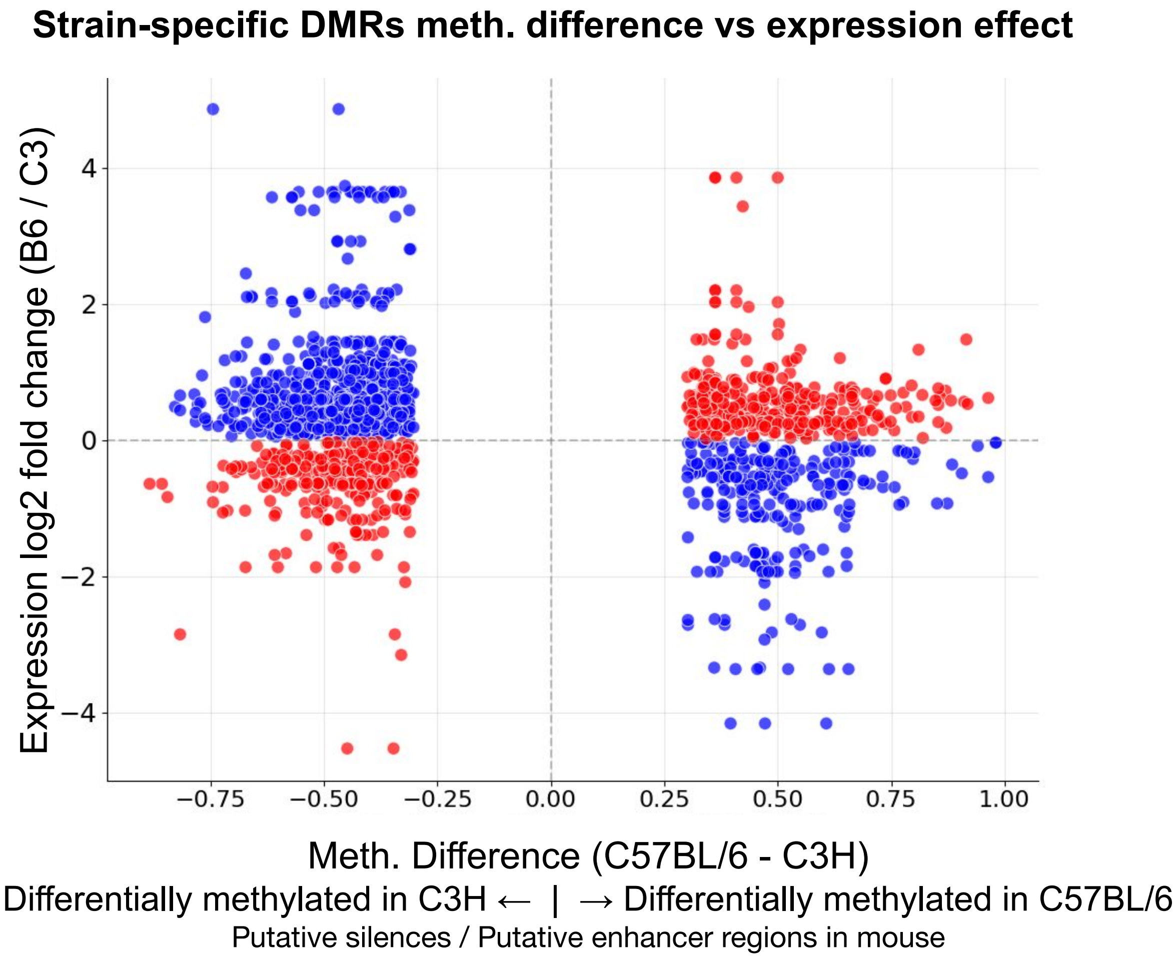


**Supplementary Figure 6. Correlation between strain-specific DNA methylation and gene expression.** Scatter plot illustrating the relationship between strain-specific DNA methylation differences (x-axis, C57BL/6 - C3H) and log2 fold change in gene expression (y-axis, C57BL/6 / C3H) for genes proximal to differentially methylated regions (DMRs). Gene-DMR pairs on the left are C57BL/6 hypomethylated (n=1073), and on the right are C57BL/6 hypermethylated (n=600). Blue points (n=1050) represent pairs with a negative correlation, consistent with an enhancer-like function where decreased methylation is associated with increased expression. Red points (n=623) denote a positive correlation, which may suggest a silencer-like function or alternative regulatory mechanism. Only gene-DMR pairs for which all DMRs near the gene are similarly methylated (i.e. all are C57BL/6 unmethylated or all are C57BL/6 methylated) are included in this analysis.

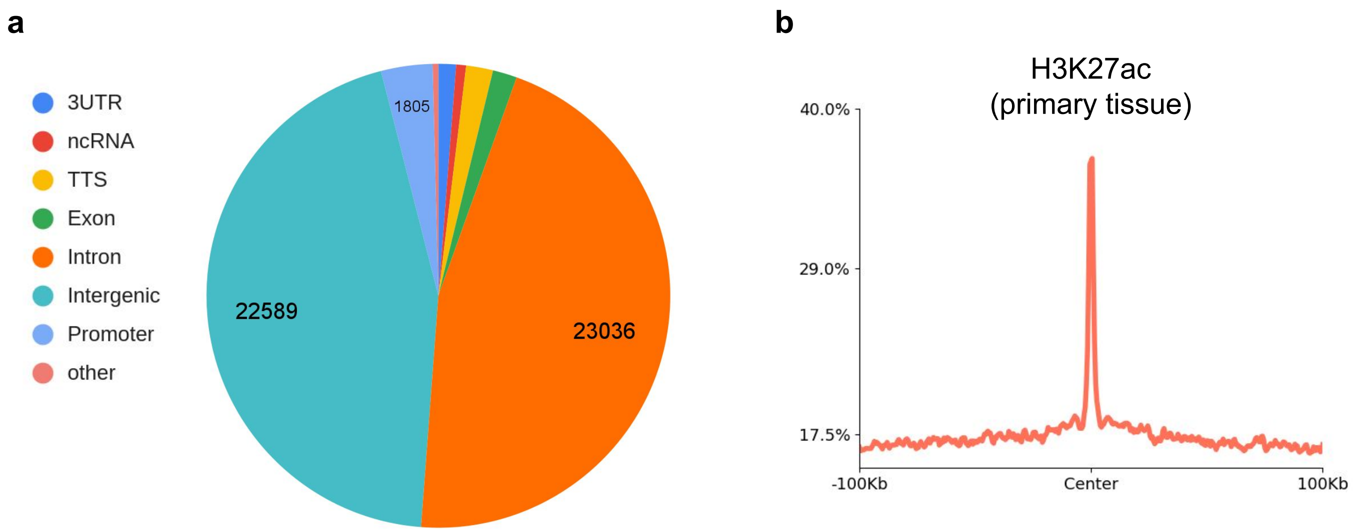


**Supplementary figure 7. Characterization of SD-ASM in humans a,** Homer genome annotations of identified SD-ASM regions. **b,** H3K27ac peaks across from multiple tissues (PMID 37770633) enrichment with SD-ASM regions.

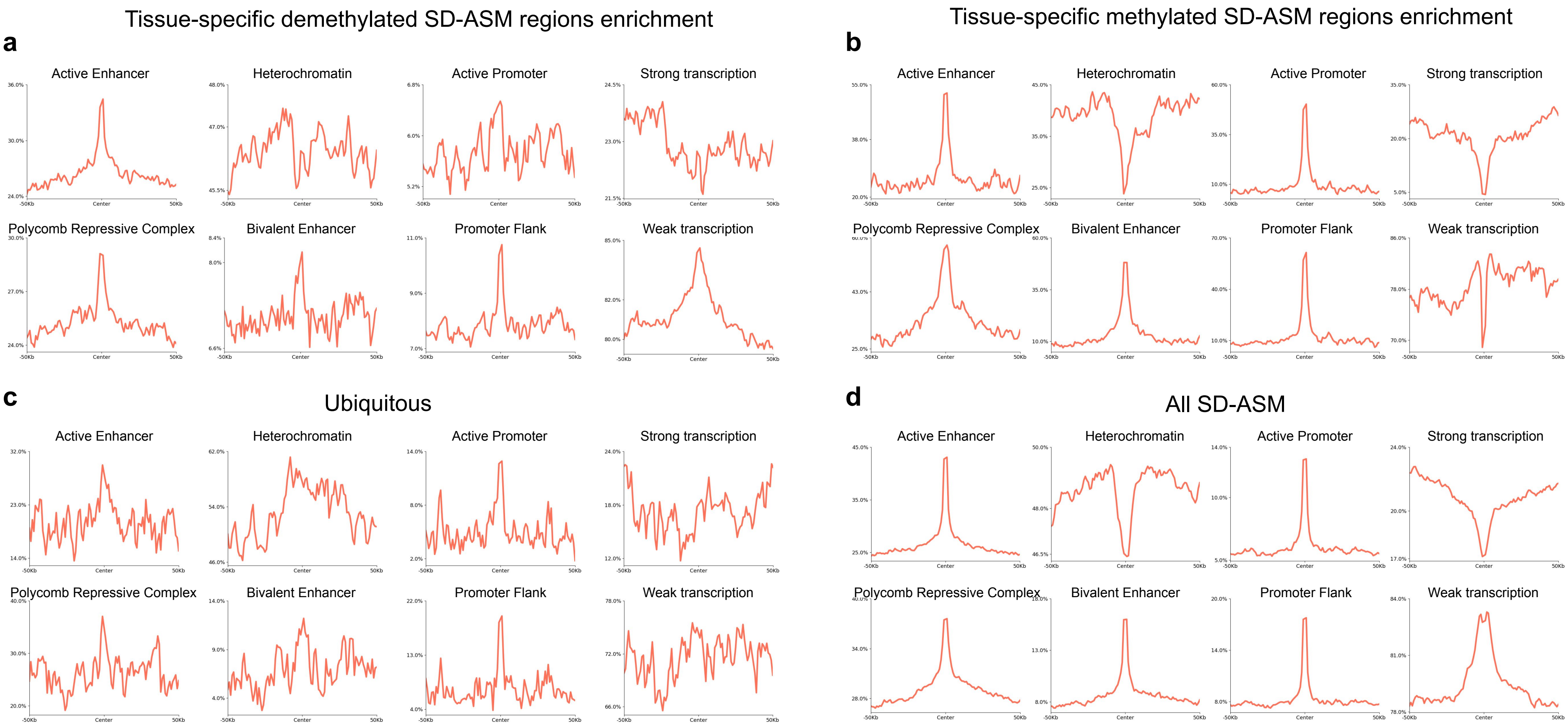


**Supplementary figure 8. Enrichment across chromHMM annotations** Enrichment with specified chromHMM (Ernst & Kellis, 2017 29120462) annotations for **a,** cell-type-specific demethylated, **b,** cell-type-specific methylated, **c,** ubiquitous, and **d,** all SD-ASM regions.

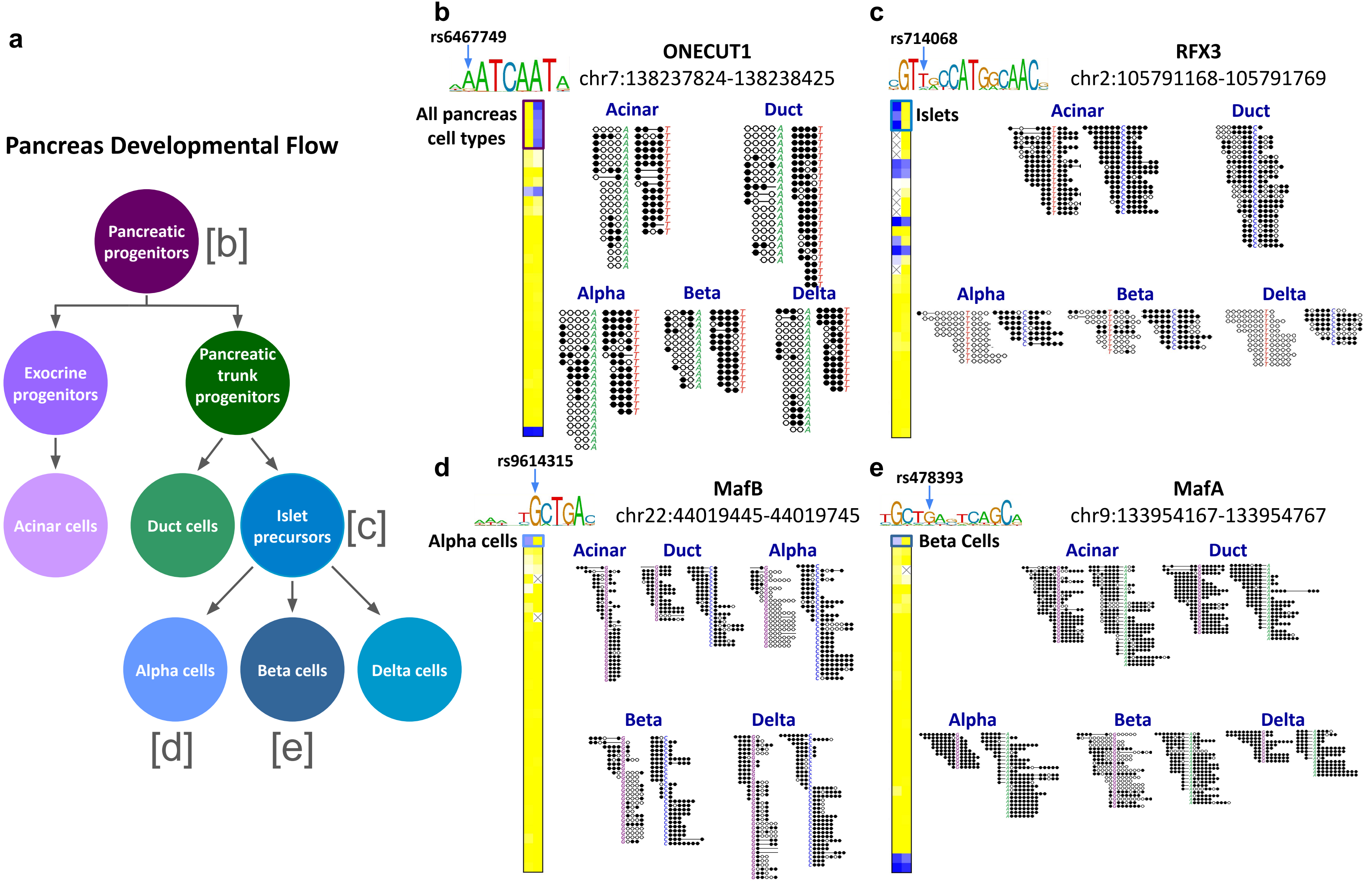
a
Pancreas Developmental Flow
Pancreatic progenitors
[b]
Exocrine progenitors
Pancreatic trunk progenitors
Acinar cells
Duct cells
Islet precursors
[c]
Alpha cells
Beta cells
Delta cells
[d]
[e]
b
rs6467749
ONECUT1
chr7:138237824-138238425
All pancreas cell types
Acinar
Duct
Alpha
Beta
Delta
c
rs714068
RFX3
chr2:105791168-105791769
Islets
Acinar
Duct
Alpha
Beta
Delta
d
rs9614315
MafB
chr22:44019445-44019745
Alpha cells
Acinar
Duct
Alpha
Beta
Delta
e
rs478393
MafA
chr9:133954167-133954767
Beta Cells
Acinar
Duct
Alpha
Beta
Delta

**Supplementary figure 9. Genetic control of DNA methylation across the pancreas developmental hierarchy. a,** Schematic representation of the pancreatic developmental flow, illustrating the lineage progression from multipotent pancreatic progenitors through exocrine and endocrine bi-progenitors to terminal cell types, including acinar, duct, and endocrine (alpha, beta, and delta) cells. Brackets indicate the specific developmental stages at which the sequence-dependent allele-specific methylation (SD-ASM) examples in panels **b–e** are established or maintained. **b,** Example of complete pancreatic SD-ASM at rs6467749 (in region chr7:138237824-138238425, hg38), where the allelic methylation difference is present across all pancreatic cell types. The SNP intersects the binding motif for ONECUT1, a transcription factor essential for early pancreatic specification. **c,** Islet-specific SD-ASM at rs714068 (in region chr2:105791168-105791769, hg38), showing allelic methylation differences restricted to the islets. This SNP overlaps the binding site for RFX3. **d,** Alpha cell-specific SD-ASM at rs91614315 (region chr22:44019445-44019745, hg38), where the genetic effect on methylation is only observed in the alpha cell lineage. The variant disrupts a binding site for the alpha cell-specific factor MafB. **e,** Beta cell-specific SD-ASM at rs478393 (chr9:133954167-133954767, hg38), demonstrating lineage-restricted methylation control. The SNP intersects the binding motif for MafA, a key regulator of mature beta cell identity and function.

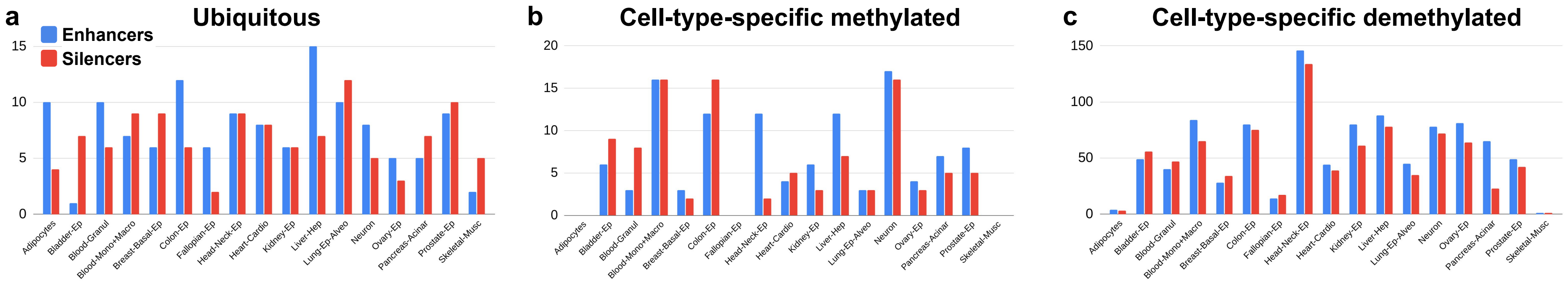


**Supplementary figure 10. Number of enhancers and silencers per cell type.** Count of the number of enhancers and silencers per cell type in **a,** ubiquitous, **b,** cell-type-specific methylated, and **c,** cell-type-specific demethylated SD-ASM loci.

**Supplementary Table 1. Liver-specific C57BL/6 unmethylated DMRs motif binding.** Shown are the enrichment statistics of the top 10 TF binding motifs from homer (n=4,439)

| TF | p-value | intersection | percent | Intersection random | percent |
|---|---|---|---|---|---|
| HNF4a | 1e-58 | 476 | 10.72% | 4412.3 | 4.73% |
| COUP-TFII(NR)/ K562-NR2F1 | 1e-46 | 994 | 22.39% | 13398.9 | 14.35% |
| EAR2 | 1e-40 | 907 | 20.43% | 12296.7 | 13.17% |
| PPARa | 1e-39 | 752 | 16.94% | 9683.8 | 10.37% |
| Foxa2 | 1e-37 | 593 | 13.36% | 7184.0 | 7.69% |
| HNF6 | 1e-37 | 405 | 9.12% | 4274.4 | 4.58% |
| FOXA1 | 1e-36 | 865 | 19.49% | 11875.1 | 12.72% |
| COUP-TFII(NR)/ Artia-Nr2f2 | 1e-34 | 1102 | 24.83% | 16295.0 | 17.45% |
| FOXA1/MCF7 | 1e-34 | 730 | 16.45% | 9715.6 | 10.41% |
| RARa | 1e-33 | 1831 | 41.25% | 30433.2 | 32.59% |

**Supplementary Table 2. Ubiquitous C57BL/6 unmethylated DMRs motif binding.** Shown are the enrichment statistics of the top 10 TF binding motifs from homer (n=193)

| TF | p-value | intersection | percent | Intersection random | percent |
|---|---|---|---|---|---|
| CTCF | 1e-7 | 16 | 8.29% | 1246.8 | 1.34% |
| BORIS | 1e-4 | 15 | 7.77% | 2059.7 | 2.22% |
| NF1 | 1e-3 | 17 | 8.81% | 3165.0 | 3.41% |
| ELF3 | 1e-3 | 28 | 14.51% | 7118.2 | 7.67% |
| ETV1 | 1e-3 | 48 | 24.87% | 14799.2 | 15.94% |
| Fli1 | 1e-2 | 41 | 21.24% | 12084.1 | 13.02% |
| Elk1 | 1e-2 | 26 | 13.47% | 6622.3 | 7.13% |
| NFAT:AP1 | 1e-2 | 8 | 4.15% | 1032.0 | 1.11% |
| EWS:ERG-fusion | 1e-2 | 24 | 12.44% | 6026.7 | 6.49% |
| Elk4 | 1e-2 | 25 | 12.95% | 6589.4 | 7.1% |

**Supplementary Table 3. Ubiquitous human SD-ASM motif binding.** Shown are the enrichment statistics of all enriched TF binding motifs from homer (n=189).

| TF | p-value | intersection | percent | Intersection random | percent |
|---|---|---|---|---|---|
| CTCF | 1e-7 | 14 | 7.41% | 939.0 | 1.12% |
| BORIS | 1e-5 | 17 | 8.99% | 1893.7 | 2.25% |
| NRF1 | 1e-4 | 16 | 8.47% | 2356.1 | 2.80% |
| NRF | 1e-3 | 18 | 9.52% | 3110.2 | 3.70% |
| Zscan4c | 1e-3 | 25 | 13.23% | 5472.3 | 6.51% |

**Supplementary Table 4. Liver-specific human SD-ASM motif binding.** Shown are the enrichment statistics of the top 10 TF binding motifs from homer (n=1155).

| TF | p-value | intersection | percent | Intersection random | percent |
|---|---|---|---|---|---|
| FOXA1/MCF7 | 1e-19 | 215 | 18.61% | 9510.9 | 9.83% |
| HNF4A | 1e-18 | 106 | 9.18% | 3288.5 | 3.40% |
| FOXA1/LNCAP | 1e-17 | 241 | 20.87% | 11409.9 | 11.79% |
| FOXM1 | 1e-17 | 215 | 18.61% | 9821 | 10.15% |
| HNF6B | 1e-16 | 170 | 14.72% | 7163 | 7.40% |
| COUP-TFII | 1e-15 | 325 | 28.14% | 17812.1 | 18.40% |
| CEBP | 1e-14 | 181 | 15.67% | 8203.1 | 8.47% |
| HNF6 | 1e-14 | 115 | 9.96% | 4265.4 | 4.41% |
| EAR2 | 1e-14 | 308 | 26.67% | 16871.3 | 17.43% |
| RARa | 1e-14 | 570 | 49.35% | 36939.3 | 38.16% |
